# Data Integration with SUMO Detects Latent Relationships Between Patients in Lower-Grade Gliomas

**DOI:** 10.1101/2020.08.10.244343

**Authors:** Karolina Sienkiewicz, Jinyu Chen, Ajay Chatrath, John T Lawson, Nathan C Sheffield, Louxin Zhang, Aakrosh Ratan

## Abstract

Joint analysis of multiple genomic data types can facilitate the discovery of complex mechanisms of biological processes and genetic diseases. We present a novel data integration framework based on non-negative matrix factorization that uses patient similarity networks. Our implementation supports continuous multi-omic datasets for molecular subtyping and handles missing data without using imputation, making it more efficient for genome-wide assays in large cohorts.

Applying our approach to gene expression, microRNA expression, and methylation data from patients with lower grade gliomas, we identify a subtype with a significantly poorer prognosis. Tumors assigned to this subtype are hypomethylated genome-wide with a gain of AP-1 occupancy in the demethylated distal enhancers. These tumors’ genomic profiles are similar to Grade IV gliomas: they are enriched for somatic chr7 gain, chr10 loss, and other molecular events that have yet to be used in the diagnosis of lower-grade gliomas as per the current WHO guidelines.

## Introduction

Biotechnologies for large-scale molecular studies of genetic diseases have advanced significantly. High throughput assays are now available to measure RNA expression, DNA methylation, and metabolite concentration in multiple tissues [1]. Given that each assay reveals a snapshot of certain cellular aspects of a disease, integrative analysis of multiple assays is often necessary for a complete understanding of its molecular etiology and important for discovering the molecular subtypes and biomarkers of the disease [2].

Molecular typing through clustering has traditionally focused on individual data types, primarily gene expression. In a few studies with multiple data types, the subtypes were generated from the different data types individually and subsequently integrated by domain experts [3–5]. Discordant results and disagreements in such analyses can be difficult to interpret and resolve. Another popular strategy is to concatenate feature matrices from multiple data types and then operate on the resulting matrix as a single data type. This approach allows the use of existing clustering techniques but requires cross-data type normalization and feature selection in individual data types before concatenation, possibly biasing the results. More sophisticated methods such as those implemented in iCluster [6], iClusterPlus [7], and Bayesian consensus clustering [8] model the probabilistic distribution of each data type and infer subtypes by maximization of the likelihood of the observed data. However, these methods require a feature selection step and make strong assumptions about the data.

The more recent methods for clustering on multi-omic data focus on similarity or distances between samples in-lieu of clustering on the feature matrices. For example, PINS [9] creates an average connectivity matrix based on the sample connectivity observed in the different data types. It then clusters using a method that depends on the level of agreement between the data types. Additionally, it perturbs the original data by adding Gaussian noise and chooses the number of clusters such that the output clusters are robust to this noise. Another popular method, Similarity Network Fusion (SNF) [10] creates a fused network of patients using a metric fusion technique and then partitions the data using spectral clustering. A more recent method, NEMO [11] calculates an average similarity matrix and then detects the clusters using spectral clustering. A comprehensive review of multi-omic and multi-view methods for the detection of subtypes is presented in Rappoport and Shamir [12].

The existing approaches have a few limitations. First, all approaches mentioned above but NEMO require that data be available for every sample and every data type, which is unlikely in most biological studies. For data with missing values, these methods need to impute missing values. But the imputation process is often computationally challenging for genome-wide analyses. Secondly, most of the methods rely on randomization to overcome computational challenges in some portion of their algorithm. Though the randomization approach can assist in finding a solution that avoids over-fitting, it also has implications for the robustness of the method. Different invocations of such a method on the same input data may produce different clustering outcomes. Lastly, statistical methods have the advantage of being able to include biological knowledge as priors. However, they often assume a parametric normal or gamma distribution of the data to make the parameter estimation tractable. Such an assumption is often not realistic and again leads to poor performance, as demonstrated in a recent comprehensive assessment of the methods for drug response prediction [13].

Here, we present a data integration framework based on non-negative matrix factorization (NMF) and showcase an implementation called SUMO (https://github.com/ratan-lab/sumo) that can integrate continuous data from multiple data types to infer molecular subtypes. SUMO handles missing data effectively and produces clusters that are robust to perturbations. Throughout the study, whenever appropriate, we compare SUMO v0.2.5 to LRAcluster v1.0 [14], MCCA v1.1 [15], NEMO v0.1 [11], PINSPlus v2.0 [9], and SNF v2.3 [10]. We use a recent benchmark [12] to show that SUMO is consistently among the best methods in identifying groups of patients with significantly differential prognosis and enrichment of clinical associations. Using simulation, we also compare SUMO to the other methods in the ability to cluster noisy datasets, to respond to perturbations, and to handle missing information.

As an application of our approach, we apply SUMO to multi-omic datasets from patients diagnosed with lower-grade glioma. Diffuse low-grade and intermediate-grade gliomas together make up the lower-grade gliomas (World Health Organization grades II and III), a diverse group of primary brain tumors with highly variable clinical behavior. Mutations in IDH, TP53, and ATRX and codeletion of chromosome arms 1p and 19q (1p/19q codeletion) have been identified as clinically relevant markers of lower-grade gliomas [16], and as of the 2016 edition of the WHO classification, gliomas are classified based not only on histopathologic appearance but also on these molecular markers [17]. Several studies have associated IDH mutations with a more favorable course of the disease, and have identified multiple subtypes with a poor clinical course [16, 18]. We identify a single cluster of patients with a significantly differential prognosis with SUMO. Patients of this cluster are enriched for genome-wide hypomethylation, somatic chr7 gain, chr10 loss, and other molecular events that have yet to be used in the diagnosis of lower-grade gliomas as per the current WHO guidelines.

## Results

### Method overview

The NMF technique aims to explain the observed data using a small number of basis components by factoring the data into the product of two non-negative matrices; one represents the basis components and the other contains mixture coefficients [19, 20]. NMF has been successfully used as a clustering method in image and pattern recognition [21–24], text-mining [25–28], and bioinformatics [29–34]. Symmetric NMF is a variant where the decomposition is done on a symmetrical matrix that contains pairwise similarity values between the data points, instead of being done directly on the data points [35]. Symmetric NMF improves clustering quality compared to the traditional formulation and forms the basis of our approach [36].

Similar to NEMO and SNF, we preprocess, transform, and standardize the data before calculating the similarity between the samples for each data type separately. If all data types are measured for all *n* samples, the similarity between samples of the *i^th^* data type form a *n* × *n* symmetric matrix *A_i_*. We then tri-factorize *A_i_* ≈ *HS_i_H^T^*, where *H* is a non-negative *n* × *r* matrix, *S_i_* is a *r × r* non-negative matrix, and *r* (≪ *n*) is the desired number of clusters. *H* in this decomposition is shared among the various data types and is a representation of the *n* samples in a *r*-dimensional subspace accounting for the adjacencies observed in all data types. Each row in *H* represents a sample, and each column in *H* denotes a cluster. If *H* is sparse, as is typically the case in NMF, a sample is assigned to the cluster corresponding to the column in which the sample has the maximum value.

Multiplicative updates are used to solve the above factorization. Since the solution is sensitive to the initial conditions, we run the solver multiple times on several subsets of samples using different initial conditions and use consensus clustering to assign the final labels and infer the optimal number of clusters (see Method section for details).

### SUMO exhibits improved performance with noisy and incomplete data

We performed several simulations to compare the accuracy of the various methods on noisy datasets with varying sample sizes and a varying fraction of missing data. We first generated a ‘ground truth’ feature matrix consisting of 200 samples and 400 features, with two distinctly separable clusters. We then simulated feature matrices of two different data types by adding different levels of Gaussian noise to this ground truth to conduct three sets of simulation experiments.

Fig S1 shows the experimental setup for the first simulation where we increase the noise in one data type while keeping a moderate amount of noise in the other data type. We generate 100 datasets for each amount of added noise and run all methods, comparing the resulting clusters to the ground truth using the adjusted Rand index (ARI). The results in Fig 1A show that all methods exhibit a median decrease in accuracy with an increase in noise. SUMO has the highest median ARI and the least variance (Fig S2) for all levels of noise.

**Figure 1:**
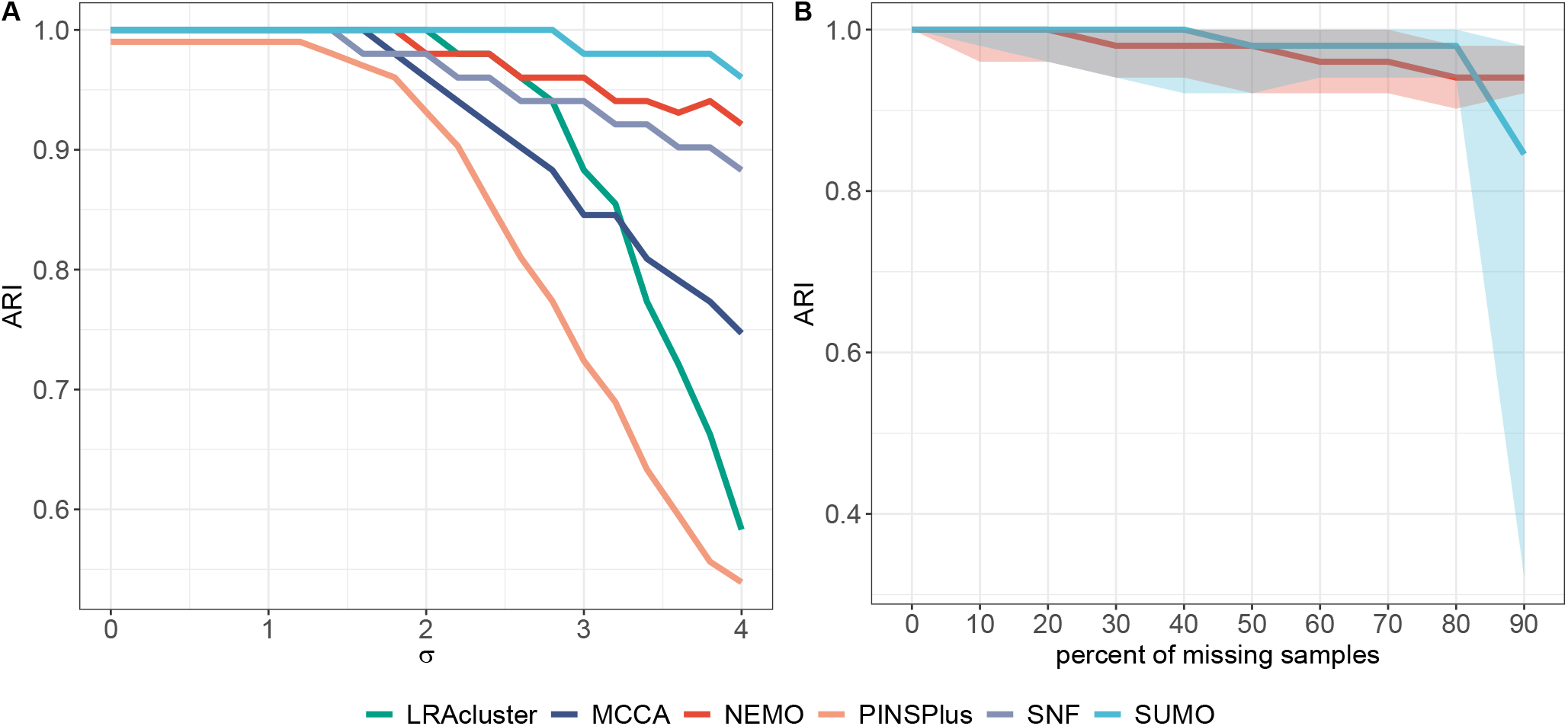
Accuracy of the six methods on noisy data and missing values. (A) Datasets were created by adding different amounts of noise to a ‘ground truth’ to simulate two distinct data types. The first data type is simulated by adding random noise from a Gaussian 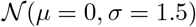 distribution, while the noise in the second data type is from a Gaussian distribution 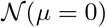 where the standard deviation is varied *σ* ∈ (0, 4)). We report the median ARI of the classification at each data point for 100 repetitions. (B) Simulated datasets were created by removing the random fraction of samples from a random data type while keeping corresponding sample data in the other data type. We plot the ARI for 100 repetitions at each data point.

Next, Fig S3 shows the experimental setup to study the impact of the sample size on the accuracy of the various tools. We again created two data types, one with a small amount of Gaussian noise, and another data type with a higher amount of Gaussian noise. Fig S4 shows the ARI of the resulting classification as an increasing fraction of samples are removed from each data type. SUMO, LRAcluster, and MCCA all score a median ARI of 1.00 for all sample sizes studied in this experiment.

Lastly, using the same setup as the second experiment, we compared SUMO to NEMO in their ability to classify samples accurately with missing data. Other methods do not handle missing data, and so were not included in this comparison. In this experiment, we removed a random fraction of samples from one data type, while preserving the data in the other data type. SUMO shows a higher median ARI compared to NEMO for most data points (Fig 1B).

### Performance of SUMO on a recent benchmark

We compared SUMO to several other methods using a recently published benchmark [12]. The benchmark consists of methylation, gene expression, and miRNA expression data from 10 cancers sequenced as part of the TCGA project. As in the original benchmark, we evaluate each method for its ability to identify a subtype that shows significantly differential survival, and is enriched for clinical annotations. We chose or calculated parameters for the methods as suggested by the authors, without considering the survival and clinical parameters that are used for assessment.

Fig. 2 depicts the performance of the various methods on the data from the different cancer types. With respect to survival, SUMO had the total best prognostic value (sum of −*log*_10_ p-values = 18.88), with MCCA being the second best with 17.48. However, the sum of p-values can be biased due to outliers, so we also counted the number of datasets for which a method’s solution obtains significantly different survival (p-value < 0.05) (Table 1). As with the original benchmark, we also evaluated if at least one of the clusters were enriched for at least one of the clinical labels. p-values for the logrank test were calculated using permutation tests [37], enrichment for discrete parameters was calculated using the *χ*^2^ test for independence, and enrichment for numeric parameters was calculated using the Kruskal-Wallis test. The p-values for clinical enrichment were corrected using Bonferroni correction.

**Figure 2:**
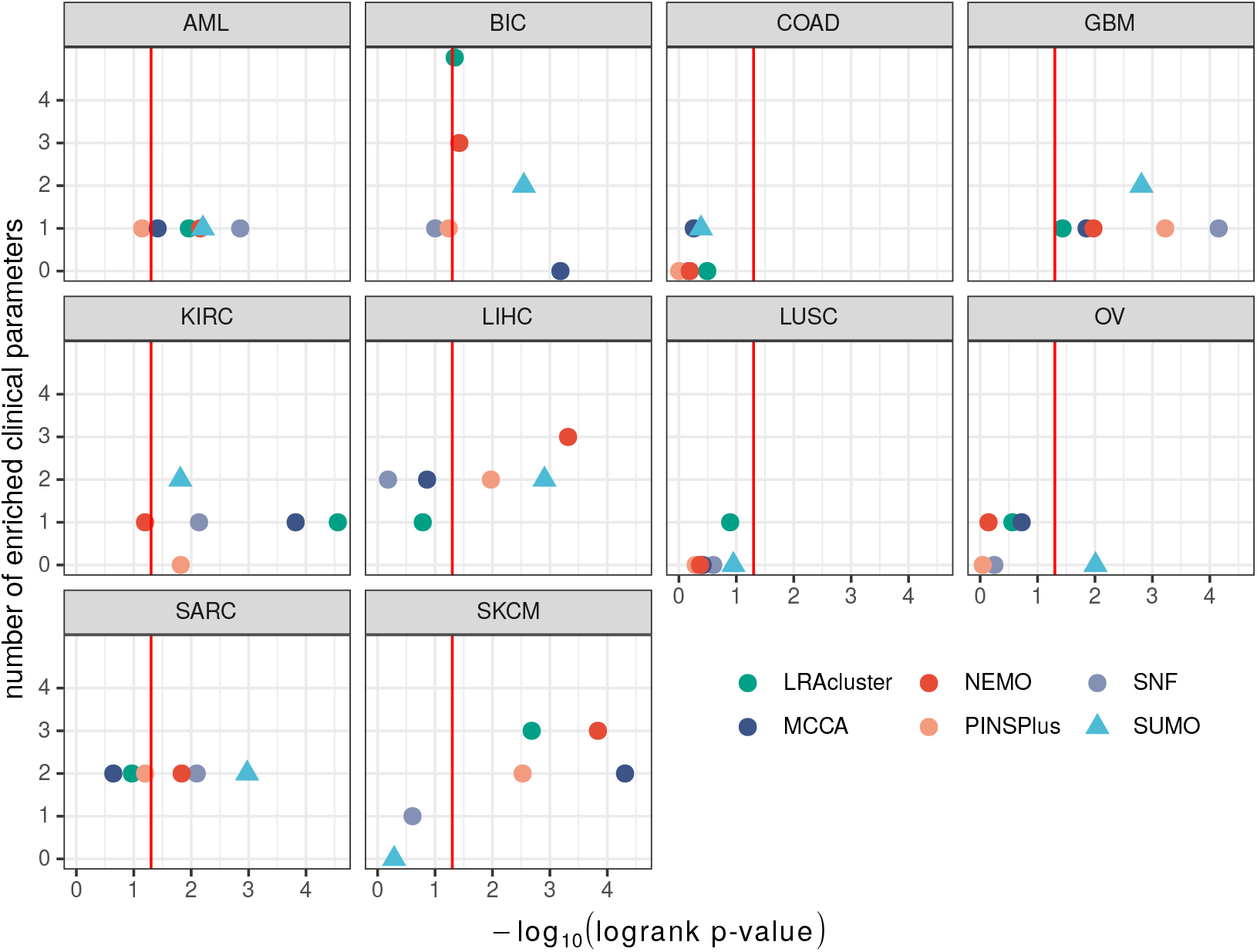
Benchmark results for the TCGA datasets. The vertical line indicates p-value= 0.05 for the logrank test, which are shown on the x-axis. The y-axis shows the number of clinical labels that were found to be enriched in at least one of the detected subtypes. SUMO results are shown using a triangle.

**Table 1:** Summary of results from the benchmark analysis. We report the number of cancers for which at least one cluster had significantly different prognosis (first column) and that had at least one enriched clinical label (second column).

| Method | Number of cancers<br>with differential survival | Number of cancers<br>with clinical enrichment |
| --- | --- | --- |
| LRAcluster | 5 | 9 |
| MCCA | 5 | 8 |
| NEMO | 6 | 8 |
| PINSPlus | 4 | 6 |
| SNF | 4 | 7 |
| **SUMO** | **7** | **7** |

Based on the results in Table 1, SUMO outperformed the other approaches, finding at least one cluster with significantly different survival in 7 out of the 10 cancers analyzed. For colorectal cancer and lung squamous cell carcinoma, none of the methods identified a subtype that showed significant differential survival. Subtypes for those cancers may be confounded of samples from a random data type while keeping corresponding sample data in the other data type. We plot the ARI for 100 repetitions at each data point. due to unknown covariates or may not exist at all, as suggested in Ma et al. [38], who found no evidence to support the existence of discrete transcriptional subtypes in colorectal cancer. SUMO is the only method to find a subgroup of patients in ovarian cancer with a significant differential survival (Fig S5A). This group of patients with poor prognosis is enriched for patients with mesenchymal tumors (Fig S5B) that are known to lead to worse outcomes [39].

All methods identified at least one cluster in Glioblastoma (GBM) with significantly differential prognosis. We used this GBM dataset to investigate the reproducibility and robustness of the methods, i.e. whether the p-values for the logrank test or the number of enriched clinical parameters would change if we changed the seed to the random number generator used by the methods and the assessment calculations. We ran each method 10 times using random seeds and found that the methods were stable to different extents on this data (Fig.S6). NEMO gave the same result in each of the 10 runs, while SUMO showed small deviations in the p-values for survival, but the remaining methods showed variation in both the p-value of the logrank test and the chi-square test used to assess the enrichment of clinical parameters. Specifically, the results for PINSPlus varied significantly in terms of survival and enrichment of clinical labels.

### SUMO analysis of TCGA-LGG identifies a cluster of patients with poor-prognosis

Several integrative approaches have been applied to understand the molecular heterogeneity and subtypes in gliomas. The largest study of diffuse grade II-III-IV gliomas to date used TumorMap [40] to integrate gene expression and DNA methylation data from around 1000 patients and identified IDH status as the primary driver of two macro-clusters [41]. The authors concluded that the IDH mutant gliomas were further composed of three coherent subgroups: (1) the Codel group, consisting of LGGs with 1p/19q codeletion; (2) the G-CIMP-low group, including gliomas without 1p/19q codeletion with relatively low genome-wide DNA methylation; and (3) the G-CIMP-high group, including gliomas without 1p/19q codeletion with higher global levels of DNA methylation. They also concluded that the IDH wild type gliomas segregated into three subgroups: (1) Classic-like, exhibiting classical gene expression signature, (2) Mesenchymal-like, enriched for mesenchymal subtype tumors, and (3) PA-like, enriched for tumors with molecular similarity to grade I pilocytic astrocytomas.

We decided to apply SUMO to subtype the lower-grade gliomas as a case study with the intent to evaluate the robustness and relevance of known glioma subtypes. We ran SUMO on the processed Level 3 gene expression, DNA methylation, and miRNA expression data for the TCGA-LGG cohort downloaded from the UCSC Xena platform [42]. We evaluated the solutions with 2 to 19 clusters according to the proportion of ambiguously clustered pairs (PAC) [43] and the cophenetic correlation [44] (See Methods for details). The PAC values suggest that the patients can be partitioned into 2 or 5 clusters, with both solutions being stable (Fig. 3A). Here, we compare our solution with 2 clusters to the findings in Ceccarelli et al. [41], and then present the solution with 5 clusters in greater detail.

**Figure 3:**
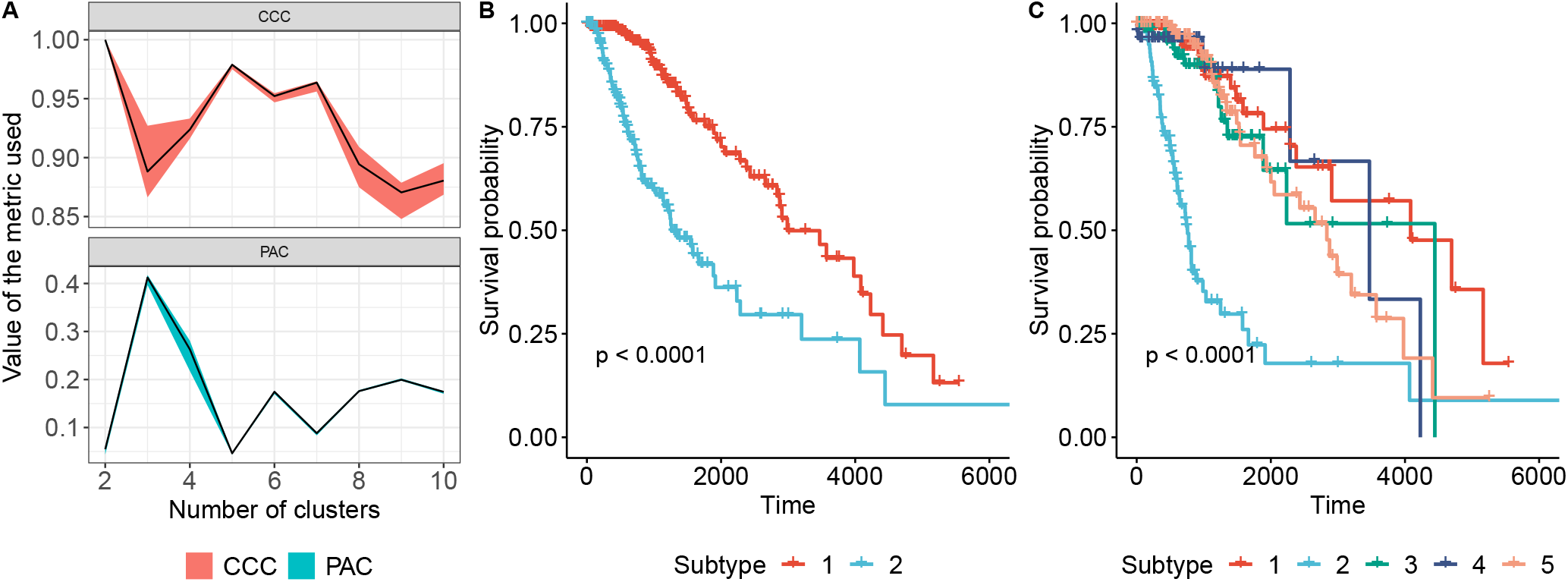
SUMO detects a single cluster showing differential prognosis in TCGA-LGG. (A) shows the two metrics used to decide the optimal number of clusters for LGG dataset. We use the proportion of ambiguously clustered pairs (PAC) (lower is better) and the cophenetic correlation (CCC) (higher is better) to select 2 and 5 as the optimal number of clusters. (B) and (C) shows the KM analysis of the subtypes detected by SUMO when 2 and 5 clusters are selected respectively.

Fig 3B shows the Kaplan-Meier survival analysis for the 2 clusters identified by SUMO. Patients assigned to the group with worse prognosis have a median survival of 1262 days compared to a median survival of 2988 days for patients assigned to the other subtype. The cluster of patients that show better prognosis include a majority of IDH mutant LGGs with 1p/19q codeletion and the majority of the IDH mutant LGG without 1p/19q codeletion with higher global levels of DNA methylation. SUMO assigns all IDH wild type patients and a subset of the IDH mutants to the subtype that exhibits a poor clinical course, and is significantly associated with higher aneuploidy (Wilcoxon rank sum test W=24243, p-value=0.0003), lower global methylation (Wilcoxon rank sum test W=58218, p-value=2.2 × 10^−16^), a higher age of diagnosis (Wilcoxon rank sum test W=24674, p-value=2.92 × 10^−5^) and a higher neoplasm grade based on histology (OR 2.45 (95% CI, 1.69 to 3.54)). Fig S7 summarizes the association of the 2 clusters with mutations, clinical phenotypes, and existing supervised classifications.

Fig 3C shows the Kaplan-Meier survival analysis for the 5 clusters as identified by SUMO. Patients assigned to Subtype 2 show a significant differential prognosis with a median survival of 758 days. Subtype 2 includes most samples (76 out of 80) that were labeled as Classic-like, Mesenchymal-like and C-GIMP low, and reported to have a poor clinical course in Ceccarelli et al. [41]. Subtype 2 also contains 18 of the 26 IDH wild type samples (labeled in Ceccarelli et al. [41] as PA-like) that were identified as having a favorable clinical course compared to other IDH wild type samples based on methylation analysis. To understand the reason for this difference, we compared the similarity between the PA-like samples assigned to Subtype 2 to (a) other samples in Subtype 2, and (b) PA-like samples assigned to other subtypes by SUMO. We determined that the PA-like samples in Subtype 2 are similar to the PA-like samples assigned to other clusters based on methylation data, consistent with Ceccarelli et al. [41]. However, the PA-like samples assigned to Subtype 2 show greater affinity to the other samples within Subtype 2 when the information from gene expression and miRNA expression were used (Fig S8).

In order to investigate if the subtypes detected by SUMO were enriched for other clinical and molecular events, we conducted enrichment analyses with the clinical phenotypes and GISTIC thresholded gene copy-number calls from UCSC Xena, along with molecular data from Ceccarelli et al. [41] and the somatic variants generated by the MC3 working group [45]. Subtype 2 is enriched for patients who are IDH wild-type and who were significantly older at the age of diagnosis (Tukey HSD test; p-value < 0.05 for all pairwise comparisons). Subtype 2 is also enriched for grade III tumors (OR 6.28 (95% CI, 3.40 to 11.59)) and significantly enriched for anaplastic Astrocytomas (p-value < 10^−5^); it is also enriched for samples with a high percentage of aneuploidy (Tukey HSD pvalues < 0.05 for all pairwise comparisons), high ESTIMATE stromal score (Tukey HSD pvalues < 0.05 for all pairwise comparisons) and high ESTIMATE combined score (Tukey HSD pvalues < 0.05 for all pairwise comparisons). This is consistent with results that suggest that the ESTIMATE scores correlate with DNA copy number-based tumor purity and high ESTIMATE scores in LGG are associated with poor outcome [46, 47].

Fig 4 summarizes some of these associations in an oncoplot. Interestingly, Subtype 2 is enriched for point mutations and amplifications of the epidermal growth factor receptor (EGFR) oncogene on Chromosome (Chr) 7. Somatic aberrations in EGFR including amplification and activating point mutations occur in ~ 57% of Grade IV gliomas but are relatively uncommon in LGGs [48]. However, 55 of the 109 patients assigned to Subtype 2 show Chr 7 gain (and hence amplification of EGFR) and Chr 10 loss, which leads to deletion of the PTEN gene, a known tumor suppressor. These chromosomal aberrations together with global hypomethylation are features unique to this subtype. As per the WHO guidelines from 2016, Chr7 gain and/or Chr10 loss are not considered in the diagnosis of Grade II/III gliomas, though other studies have suggested that these events are clinically relevant, and their inclusion in the diagnostic criterion could lead to the reclassification of several LGGs into GBMs [49].

**Figure 4:**
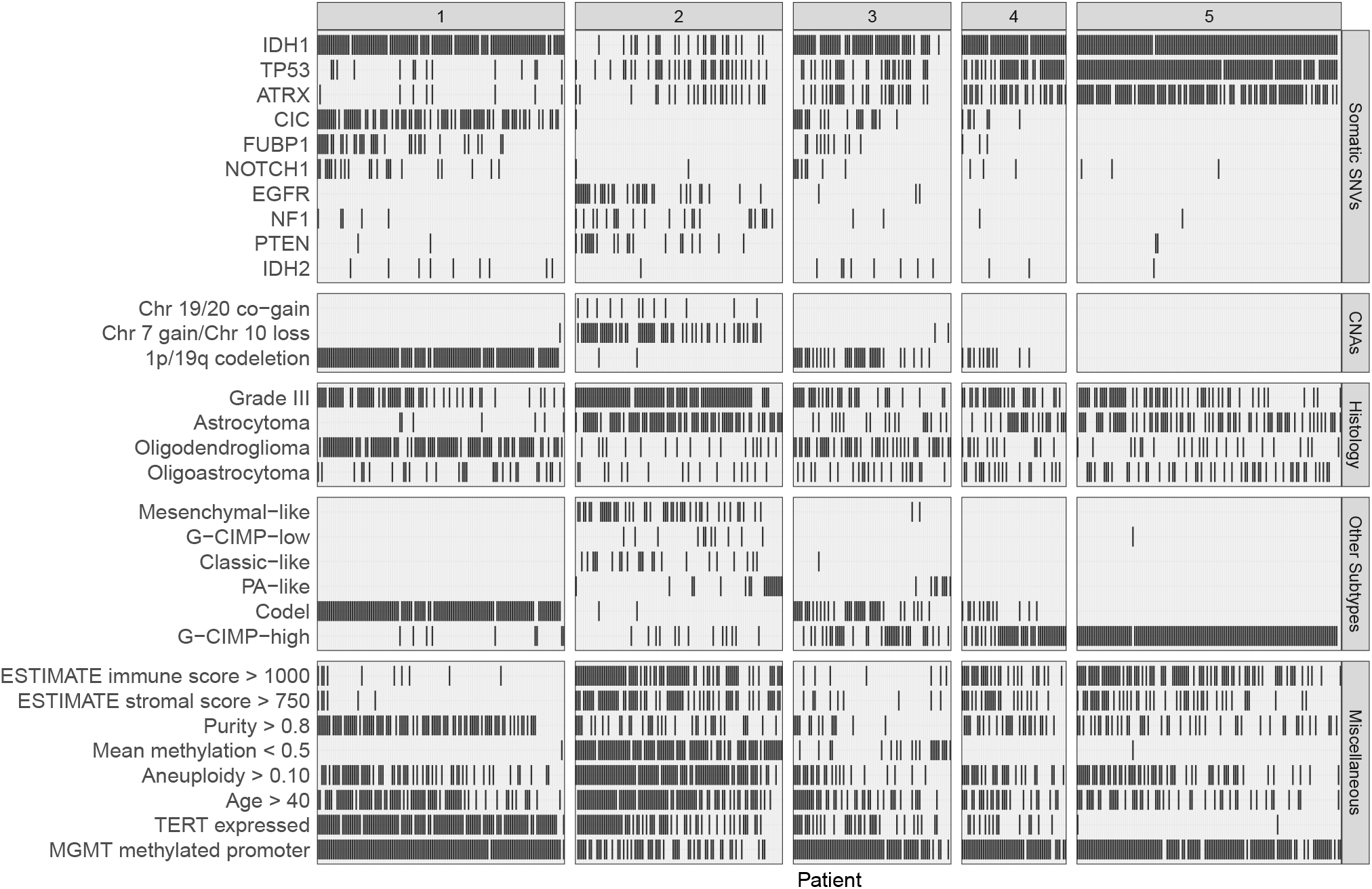
Oncoplot showing enrichment of molecular and clinical features in the various subtypes.

Since tumors are a complex milieu of numerous cell types, we hypothesized that the microenvironment plays an important role in the determination of these subtypes. To investigate this, we downloaded the xCell scores corresponding to enrichment of 64 different immune and stromal cell types in these TCGA samples [50]. Hierarchical clustering of the mean enrichment scores for the various cell types in Fig 5A shows that the cellular profile of Subtype 2 tumors is more similar to GBMs than to the other LGGs. More importantly, astrocytomas assigned to Subtype 2 have higher enrichment scores for astrocytes, similar to those calculated for GBM samples, and significantly higher than astrocytomas assigned to the other subtypes (Fig 5B). xCell scores are calculated using gene expression, but we observe similar results on analysis of methylation data using MIRA [51]. Subtype 2 samples show lower methylation and higher regulatory activity at astrocyte-specific elements (Fig 5C) compared to the other subtypes. These differences in cellular population also manifest in principal component analysis of gene expression and methylation data when we consider the LGG and GBM samples together (GBMLGG dataset from UCSC Xena). In PCA analyses of expression and methylation (Fig S9), the first principal component shows the similarities between Subtype 2 and the GBM samples. These findings along with the observed chromosomal aberrations suggest that LGGs assigned to Subtype 2 should be treated more aggressively and potentially reclassified as GBM.

**Figure 5:**
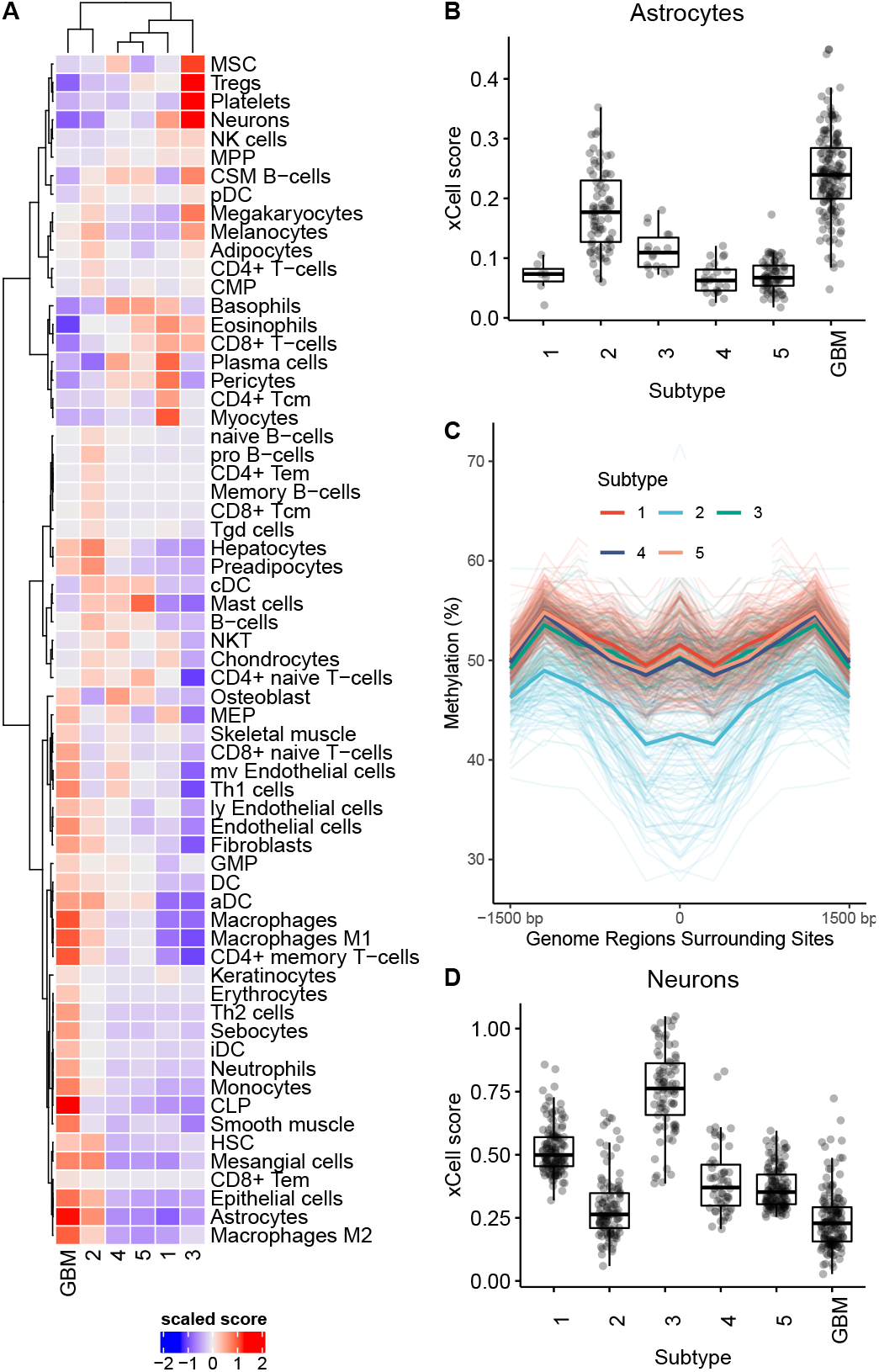
Subtype 2 shows similarities to GBM, and Subtype 3 is enriched for neurons. (A) is a heatmap that shows the mean xCell enrichment scores for the LGG subtypes and GBM corresponding to 64 cell types, with Subtype 2 and GBM sharing enrichment of several cellular populations. (B) Astrocytomas assigned to Subtype 2 show higher xCell scores compared to astrocytomas that are assigned to the other LGG subtypes. (C) Tumors assigned to Subtype 2 show lower methylation and higher regulatory activity at astrocyte-specific elements. The mean methylation levels are shown using dark line. (D) Tumors in Subtype 3 are enriched for neuronal cells.

Our enrichment analyses show that global hypomethylation is a hallmark of Subtype 2 tumors. In order to investigate this further, we used ELMER [52] in an unsupervised mode to compare Subtype 2 tumors to the other LGGs. ELMER identified 16,822 distal probes that were hypomethylated in Subtype 2 samples (adjusted p-value < 0.01 and methylation difference between means of the groups > 0.3). For 382 of those probes, their methylation status was inversely proportional to the expression of a putative target gene. These target genes are enriched for biological processes such as extracellular matrix (ECM) organization and molecular functions such as kinase binding (Fig 6A). ECM is known to be an important determinant of glioma invasion and kinase binding is activated in gliomas [53–55]. Fig 6B shows the motifs that are enriched around the 382 probes that are identified as putative distal enhancers. The motifs that show the highest enrichment correspond to the Fos and Jun transcription factor gene families. Fos genes encode leucine zipper proteins that can dimerize with proteins of the JUN family, thereby forming the early response transcription factor complex AP-1. As such, the FOS proteins have been implicated as regulators of cell proliferation, differentiation, and transformation [56]. More specifically, we find that the expression of FOSL1, which contributes to regulation of placental development is significantly higher in Subtype 2 tumors, and higher expression of the gene is associated with worse prognosis [57]. These results are in agreement with other published studies that show that AP-1 binds to demethylated regions in G-CIMP-low tumors, but we find this to be true for all samples assigned to Subtype 2 [58].

**Figure 6:**
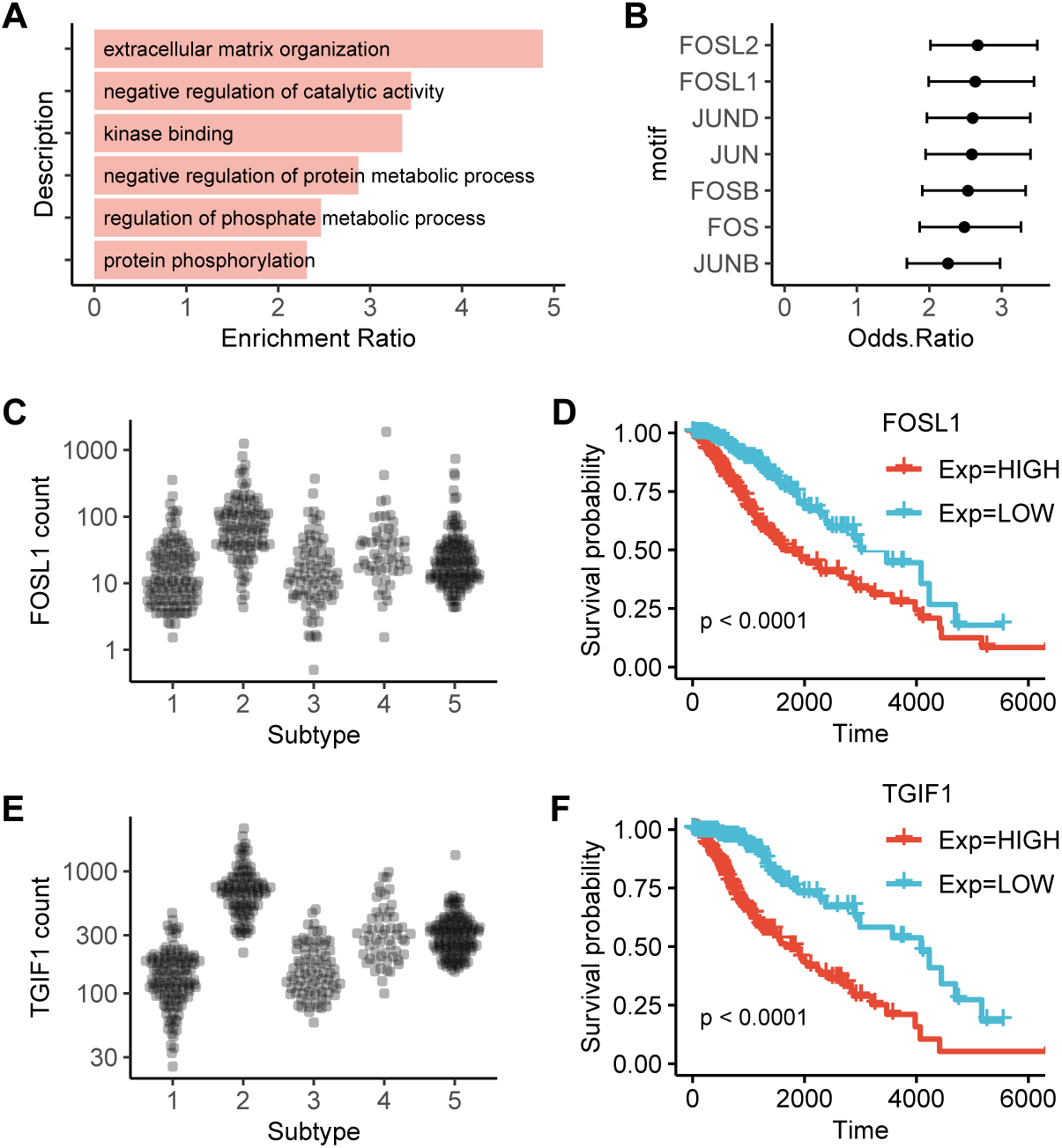
Hypomethylation in Subtype 2. (A) Enrichment of molecular function and biological processes in genes which are regulated via hypomethylation of distal enhancers in Subtype 2. (B) Motif enrichment analysis of regions around putative distal enhancers probes that regulate expression and are hypomethylated in Subtype 2 tumors. We show the 95% confidence interval of the odds ratio, and only show motifs which have a lower odds ration > 1.5. FOSL1 occupied enhancer elements show significant hypomethylation in Subtype 2 tumors. FOSL1 gene is upregulated in Subtype 2 (C), and higher expression is associated with worse survival (D). TGIF1 expression is highly correlated to enhancer hypomethylation. TGIF1 is upregulated in Subtype 2 and higher expression is associated with worse survival.

Since members of a TF family have very similar DNA binding domains, it is challenging to identify the TF that binds *in-vivo* to a region containing a motif. But we instead searched for cases where the motif occupancy of hypomethylated enhancers accompanied an increase in expression for at least one member of that TF family. Furthermore, we checked to see if the expression of the TF was significantly correlated with survival (logrank test, p-value < 0.00001). We again found an enrichment of AP-1 containing enhancers, which is a common feature of many cancer types. Interestingly, we found that TGIF1 expression was highly correlated with the degree of enhancer hypomethylation even for motifs where we did not expect TGIF1 to bind. Fig 6E and Fig 6F show that expression of TGIF1 is higher for Subtype 2 tumors and higher expression of the gene is predictive of worse prognosis. It is possible that these correlations are due to indirect effects caused by TF networks. TGIF1 is involved in regulation of cell development and maturation, and other studies have included TGIF1 in prognostic gene sets for Glioblastoma though the role of TGIF1 in gliomas is not clear [59].

We also find significant enrichment of clinical and molecular features in other subtypes. Subtype 1 is enriched for Oligodendrogliomas (p-value < 1.0 × 10^−5^), mutations in the TERT promoter and high expression of TERT (Tukey HSD test; p-value < 0.05 for all pairwise comparisons), high tumor purity (Tukey HSD test; p-value < 0.05 for all pairwise comparisons), 1p/9q co-deletion, and mutations in CIC, a known tumor suppressor. 128 of the 130 patients in Subtype 1 have a methylated promoter for MGMT (post hoc test of residuals for *χ*^2^ test, p-value: < 1.0 × 10^−5^). MGMT promoter methylation is associated with better response to alkylating chemotherapy, suggesting that patients assigned to Subtype 1 are more likely to respond to temozolomide [60].

Subtype 3 is enriched for the neural (NE) subtype detected in previous gene-expression studies [61]. The NE subtype has previously been related to the tumor margin where increased normal neural tissue is likely to be detected [62]. Consistent with this hypothesis, we find that the tumors assigned to Subtype 3 have lower tumor purity (Tukey HSD test; p-value < 0.05 for all pairwise comparisons except with Subtype 5) and high enrichment score for neurons (Fig 5C. Subtype 4 and Subtype 5 are both enriched for G-CIMP high samples, although Subtype 5 is enriched for mutations in ATRX (post hoc test of residuals for *χ*^2^ test, p-value: < 10^−5^), and shows a higher enrichment for Mast cells which are known to induce release of selective inflammatory cytokines such as IL-4 with anti-glioma activity leading to improved prognosis [63].

## Discussion

We present an approach to integrate multi-omic data and use it to subtype LGG through the integration of gene expression, DNA methylation and miRNA expression data. Our method is based on symmetric NMF and can be easily extended for various applications. For instance, we develop an implementation, SUMO, for unsupervised learning by regularization of the cluster indicator matrix using the Frobenius norm. It can be modified into a semi-supervised learning to classify samples after the inclusion of priors based on a phenotype of interest, as suggested in other studies [64, 65]. Additionally, we find that the primary LGG tumors show a significant difference in survival based on histological grade within existing subtypes (Fig S11). Such a semi-supervised framework will allow for integration of clinical observations with molecular information.

SUMO improves on existing methods in its ability to handle noisy and missing data. We compared SUMO to several existing methods for integrative clustering. SUMO produces consistently reproducible results on a recently published benchmark. The benchmark uses differential survival and enrichment of a small number of clinical labels in the resulting clusters as metrics for assessment of subtyping methods. However, it is important to remember that subtypes of a disease that are biologically different can lead to similar survival. For example, we find that PA-like samples from Ceccarelli et al. [41] get classified by SUMO primarily into two groups based on gene expression and miRNA-expression, even though the two groups are not significantly different in terms of survival. SUMO focuses on the integration of continuous data types such as expression, methylation, and metabolomics. Sparse and noisy data types such as somatic mutations can be included for integration after limiting the features to those that have a known role in the disease. Alternatively, such data types can be converted into continuous data types by use of network propagation techniques and then included as input to SUMO [66].

We applied SUMO for the detection of subtypes in lower-grade gliomas, and identified a single subtype with differential prognosis compared to the other subtypes. We show that this subtype includes all previously studied groups of patients with features that are associated with a poor outcome. Like GBM, gain of chr7, loss of chr10 and global hypomethylation appear to be hallmarks of this subtype, and our analyses suggest that LGGs assigned to Subtype 2 should be treated more aggressively and potentially reclassified as GBM. This subtype should also be analyzed separately in clinical trials as its molecular differences may make it susceptible to different drugs with respect to the other LGG subtypes. It is also an open question as to whether or not this subtype regrows/recurs faster after neurosurgical resection compared to the other subtypes. Additionally, we also found that the hypomethylated distal enhancers in this subtype are enriched for AP-1 binding. This has been shown to be a feature of G-CIMP-low tumors, but we find it to be characteristic of most Subtype 2 tumors. We also identified TGIF1 expression to be inversely proportional to the global hypomethylation, and predictive of prognosis, even though its role in glioma is not clear.

A common post hoc analysis to molecular subtyping is identification of feature or sets of features that can be used as markers or surrogates for the various subtypes. SUMO includes a mode to build a tree-based model that can predict the importance of each feature for each of the detected subtypes. For example, we identified an clinically relevant subtype of LGG with differential prognosis compared to the other subtypes. According to our analysis, the non-CpG island methylation probes in the proximity to the gene CLCF1 are the best marker for the subtype. Fig S12 shows the beta values of the samples for the three methylation probes that have the highest explanatory values for the classifier.

In summary, SUMO is as a molecular subtyping method that can handle noise and missing data that commonly exist in genomic datasets and can be extended for other applications. Our study suggests that NMF-based multi-omic integration is a promising approach that can be applied to a wide range of biomedical datasets and can provide valuable biological insight.

## Methods

Our approach is based on non-negative matrix factorization, where the factorization is jointly performed on the similarity matrices calculated for all data types separately. After removal of outliers and data normalization, we first transform the feature matrix from the *i^th^* data type into a similarity matrix *A_i_* between the samples and then tri-factorize *A_i_* into *HS_i_H^T^*, where *H* is the cluster indicator matrix that is shared across all data types. The objective function used for computing the tri-factorization accounts for the missing samples and the difference in sample size for the various assays. Lastly, to produce a robust clustering, we run the solver multiple times and apply consensus clustering to obtain the final clusters. Now, we describe these steps in details.

### Data preprocessing

Data preprocessing involves (a) filtration, (b) transformation, and (c) normalization of each data type separately. The filtering process removes features that are not informative; for example, we removed genes that had zero counts in most samples. Even though our approach can handle missing values, removing features and samples with a large fraction of missing values (> 10%) often speeds up computation and improves the classification if it does not remove a significant fraction of samples.

The transformation process is data-dependent. We use a variance-stabilizing transform to convert abundance in count datasets, for example as in RNA-seq, to yield a matrix of values that are approximately homoscedastic (with constant variance along the range of mean values). This had an additional advantage of reducing the effect of outliers in the dataset. We use M-values over beta values to transform methylation datasets [67]. If batch information is known, we use ComBat [68] to adjust for batch effects in this step.

In the normalization step, we perform feature standardization to make the value of each feature in the data have zero-mean and unit variance.

### The construction of similarity networks and matrices

Let *n* be the number of patient samples *s*, that are found in the dataset of every data type and let *t* be the number of data types e.g., gene expression or DNA methylation. In this step, we construct a similarity network *N*, which we represent as a set of *n* × *n* similarity matrices {*A*_1_, *A*_2_, · · ·, *A_t_*}, where *A_k_*(*i, j*) = (*a_ij_*(*k*)) and *k* is used as an index for the data type. *a_ij_*(*k*) represents the similarity between two samples *s_i_* and *s_j_* calculated from the features of the *k*^th^ data type, *k* = 1, · · ·, *t*.

For each data type *k*, we assume its data is represented in a matrix (*f_ij_*) containing *n* sample rows and *p* feature columns. We calculate *A_k_* as a radial basis function of the Euclidean distance 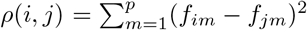 between the samples *x_i_* and *x_j_*:

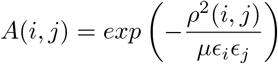

where *μ* is a hyperparameter with a default value of 0.5 and *ε_i_* represents the average distance between *x_i_* and its *K* nearest neighbors:

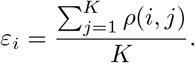

We set the number of nearest neighbors *K* equal to 10% of the samples in the data type. The selection of this parameter can effect the results, and we recommend setting it to 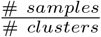 if the number of clusters is known.

The Euclidean distance is appropriate for normalized count datasets, such as those that arise from gene expression or DNA methylation data. However, depending on the data type and the application, different distances or similarity metrics may better represent sample relationships. For example, cosine similarity has been shown to be a better metric for calculation of similarity between single cells in the single-cell sequencing for transposase accessible chromatin (scATAC-seq) [69].

### Joint tri-factorization of the similarity matrices

Each matrix *A_i_* of the multiplex network *N* is symmetric and non-negative. We tri-factorize *A*_1_, *A*_2_, · · ·, *A_t_* as follows:

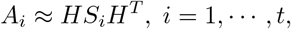

in which *H* is a *n* × *r* matrix shared across the data types and *r* is the desired number of clusters such that *r* ≪ *n* (Fig S10B).

We compute the above tri-factorization by minimizing the following objective function:

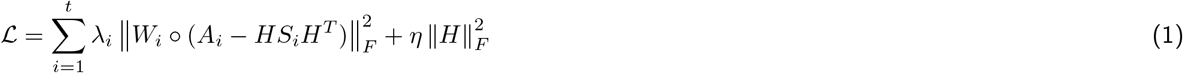

where ◦ denotes entry-wise multiplication for matrices, and *H* and *S_i_* are both constrained to be non-negative. The first term of the objective function measures the divergences between *A_i_* and *HS_i_H^T^* using the Frobenius norm in each data type. For each data type, measurements may be not available for all the *n* samples, thus leading to missing entries in the matrix *A_i_*. We use *W_i_* to remove the missing values, where

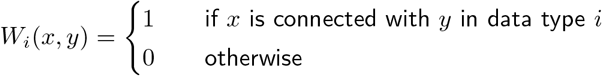

Then we add an another factor 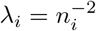 to account for the imbalance in the number of entries among *A_i_*(*i* = 1, …, *t*), where *n_i_* is the number of samples for the *i*^th^ data type.

The second term of the objective function is used to enforce sparsity on the matrix *H*, hopefully leading to a non-overfitted result and the hyperparameter *η* is used to balance the contribution of these two terms.

Note that the cost function in Eqn. 1 is convex in either but not both *H* and *S_i_*. The following multiplicative updates are used to solve the optimization problem given in Eqn. 1 [70].

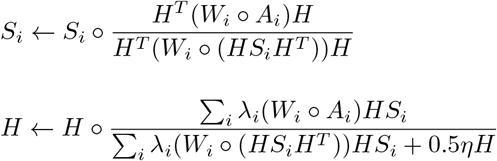

As the algorithm iterates using the updates, *H* and *S_i_* converge to a local minimum of the cost function. We apply above rules iteratively while alternating fixed matrices, keeping track of objective function value 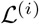 until it satisfies

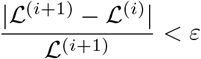

where *ε* is a predefined threshold, or the maximum number of allowed iterations are reached.

Since the solution is relatively sparse, we can assign each sample (represented by a row in *H*) to the cluster corresponding to the column that contains the maximum value, as depicted in Fig S10C. In practice, the solution can be sensitive to the initial conditions. We discuss the details of this in the implementation details, but briefly, we run the above solver multiple times and then use consensus clustering to get the final assignments.

#### Derivation of multiplicative-update rules

For the objective function Eqn. 1, when we update matrix *S_i_*, matrices *H* and *S_j_* (*j* ≠ *i*) should be fixed, thus it would be an optimization problem about the matrix *S_i_*, that is,

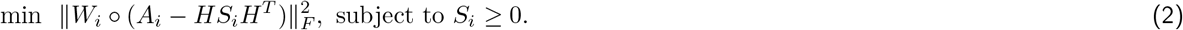

The corresponding Lagrange function of Eq. (2) is

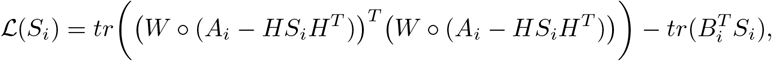

where *B_i_* ≥ 0 is the Lagrange multiplier for *S_i_*, and *tr*(·) represent the trace of matrix *X*. Then

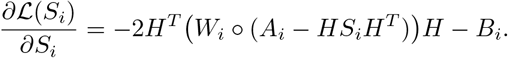

Let 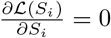, thus

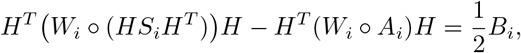

and

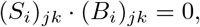

thus *S_i_* satisfies

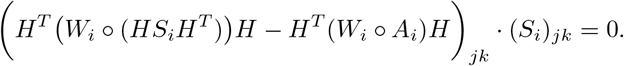

We obtain the update formula for *S_i_* as follows:

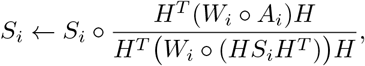

where ◦ and ÷ denote entry-wise multiplication and division for matrices, respectively. Similarly, when we update matrix *H*,

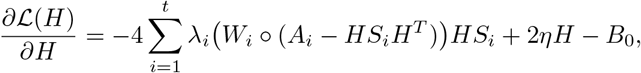

where *B*_0_ ≥ 0 is the Lagrange multiplier for *H*. Thus, *H* satisfies the following equations:

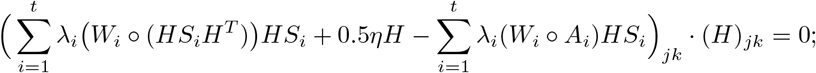

Then, we obtain the following update formulas for *H*:

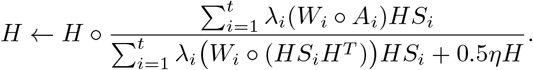

### Implementation details

SUMO (https://github.com/ratan-lab/sumo) is specifically designed to integrate multi-omic data for molecular subtyping. It consists of four subroutines. It allows the user to construct the multiplex network from normalized feature matrices (*sumo prepare*), tri-factorize the multiplex network to assign samples to the desired number of clusters (*sumo run*), compares the assignments to another classification using multiple metrics (*sumo evaluate*), and detect the importance of each feature towards each cluster (*sumo interpret*), which facilitate the discovery of biomarkers and molecular signatures.

SUMO is available in the form of a command-line tool on GitHub (https://github.com/ratan-lab/sumo) and at The Python Package Index (https://pypi.org/project/python-sumo/).

#### Support for missing data

Biomedical studies measure a large number of molecular parameters. Almost every dataset has missing entries. Most methods for molecular subtyping require perfect data. This implies that that both samples and features that have missing entries have to be removed or the missing entries are imputed in the pre-processing stage. SUMO takes a different approach. It scales the calculated distance between a pair of samples by the number of common features available for both samples. If sufficient overlap (by default at least 10% of features) is not found, the distance is set to *NA* (not available). A missing value in an adjacent matrix *A_i_* is equivalent to a missing edge between two nodes in the multiplex network and is masked during factorization as we describe in the last section.

#### Consensus clustering

As we mention in the last section, our iterative solution using multiplicative rules is sensitive to the initial conditions. Both initialization and convergence speeds are important factors to consider when formulating the appropriate factorization algorithms [71]. Our method utilizes an SVD based initialization approach to set the initial *H* to be the average similarity matrix across all data types. This method reduces residual error and provides faster convergence than using random initialization. However, we still have to set *S_i_* randomly; as such, the algorithm does not guarantee convergence to a local minimum. Here, we set the diagonal entries of each *S_i_* to be absolute singular values, that are derived from the SVD decomposition of the corresponding *A_i_* matrix. We repeat the factorization *n* times, each time including 95% of the total samples in calculating the cluster assignments from *H* and a residual error *RE_i_* for that run. We create a consensus matrix from these *n* assignments that is weighted to incorporate the residual error (RE) of each factorization in a dataset with *t* data type as follows.

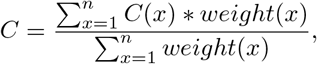

where

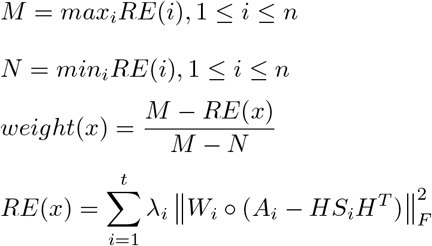

We use the Normalized Cut clustering algorithm [72] on this consensus matrix to assign the final cluster labels.

#### Estimating the optimal number of clusters

Estimation of an optimal rank for NMF is a challenging problem. It is common to compare several solutions based on a clustering metric [73]. We implement two popular metrics that leverage the consensus matrix to help the user in the determination of stable solutions to the factorization. The first metric is the cophenetic correlation coefficient (CCC) [74]. It measures the Pearson correlation between sample distances and its hierarchical clustering. A higher CCC value is considered better. The second metric is the proportion of ambiguously clustered pairs (PAC), which is defined as the proportion of the consensus matrix values in (0.1, 0.9) range. Based on our experiments, we recommend investigating factorization rank values for which the PAC score is less than 0.1, and the CCC value is high (typically > 0.95). Increasing the number of repetitions of the solver can assist in identification of the optimal number of clusters, but as we show in Fig S13 using the acute myeloid leukemia (AML) dataset from benchmark data [12], we can identify one of the stable solutions in a small number of repetitions. Similarly, we use the same dataset to show in Fig S14 that the trends observed in the PAC curve and the CCC curve are preserved for a wide range of values corresponding to the number of samples that are removed in each iteration [0,0.1]. In the current default setting, we run 60 repetitions of the solver. With each run we randomly remove 5% of the samples, while making sure that each sample will be clustered at least once. We then use random subsets of 50 runs to create multiple weighted consensus matrices as described in previous section. While only one of the matrices is utilized to call sample labels, the CCC and PAC metrics are calculated for every one of them, providing a robust assessment of stability of factorization results.

#### Identification of biomarkers

Once the subtypes are assigned, a frequent challenge is to identify a set of features that correlate with the cluster separation. These can be used as markers for the assignment of future samples and can aid in understanding the differences between the groups. To this end, we first train a gradient boosting classifier implemented in LightGBM [75]. We use 80% of the features for training this model while performing hyperparameter optimization of the model using a random search with 5-fold cross-validation to avoid overfitting. When we have this model, we calculate the Shapley values of all features for each identified cluster. The features with a Shapley value greater than 1 are considered to be important in driving separation of that cluster.

## Acknowledgments

This work was supported by NIH award (P30CA044579) to the UVA Cancer Center, and by NIGMS award GM128636 (NCS). JY Chen and LX Zhang were partially supported by the Singapore Research Foundation (Grant No. NRF2016NRF-NSFC001–026). JTL was supported by NIH training grant (NLM; 5T32LM012416) and the UVA Cancer Center.

## Supporting information

**Figure S1:**
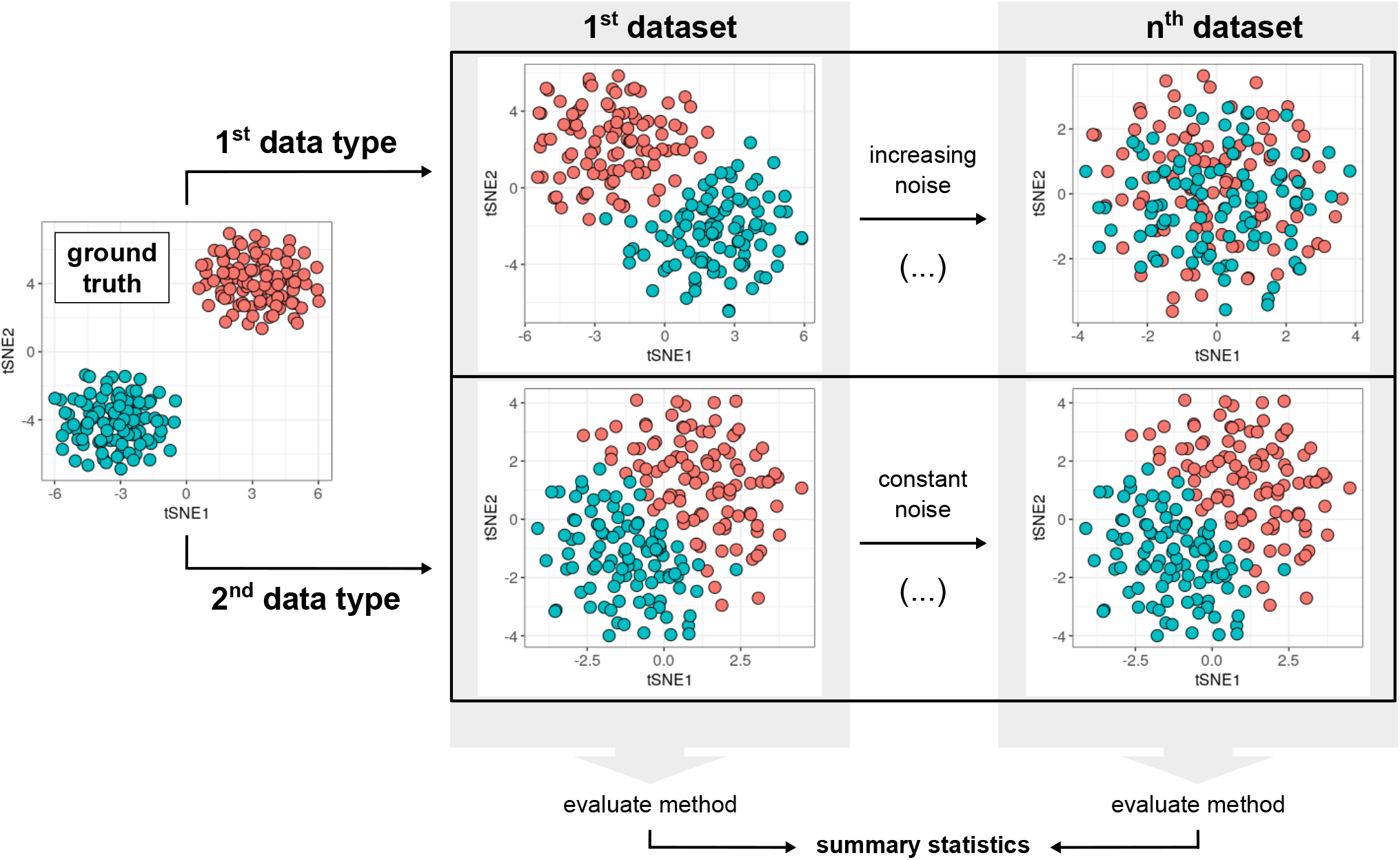
Experimental setup to compare the accuracy of the various methods on noisy data. During simulation, noise in one of the data types remains constant (generated from Gaussian distribution 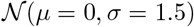). Another data type contains data with increasing noise, randomly generated from the Gaussian distribution as a function of standard deviation, and constant mean. Here we show two sampling points for following sets of parameters (*μ, σ*) ∈ {(0, 1), (0, 4)}.

**Figure S2:**
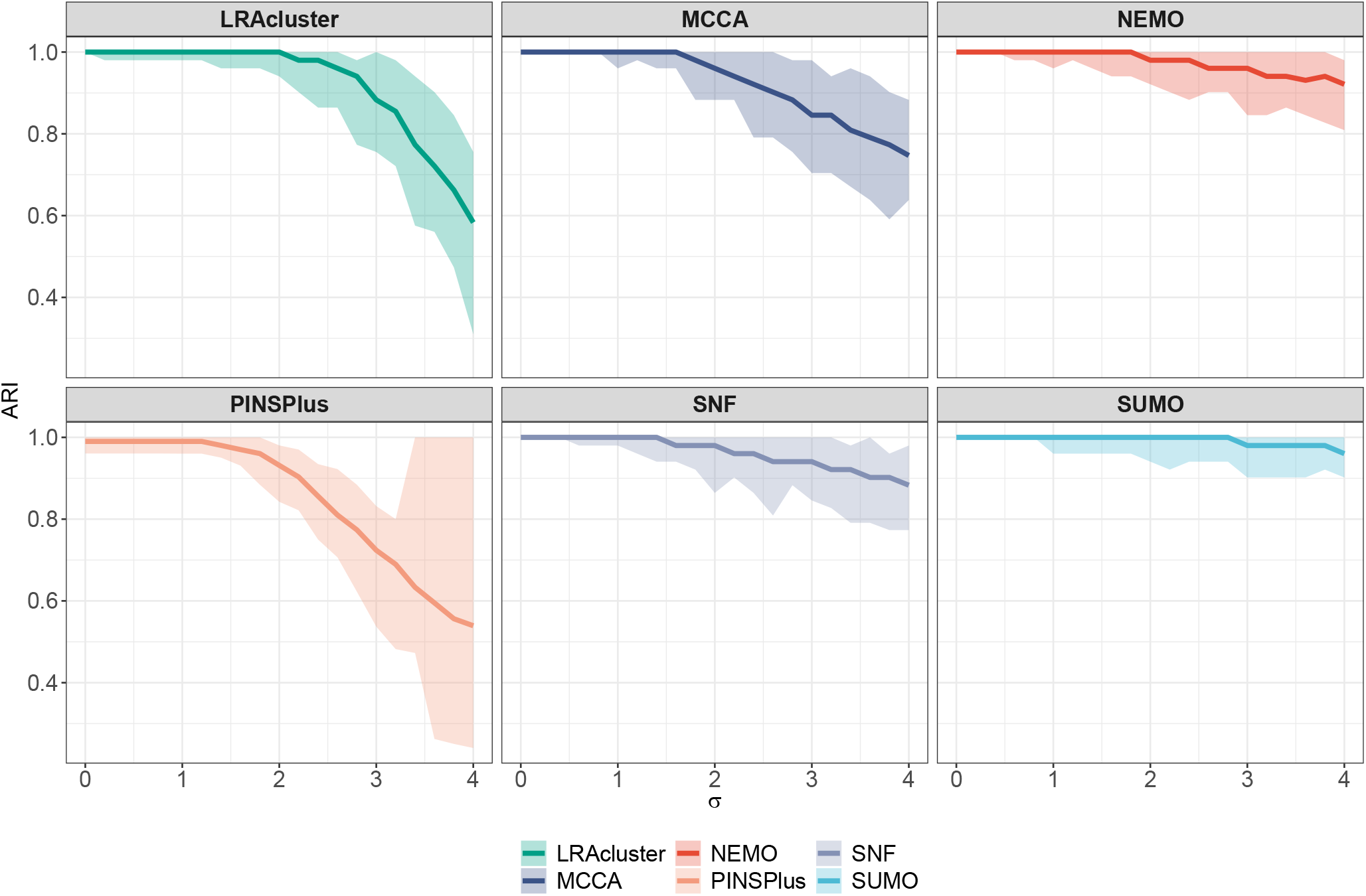
Adjusted Rand Index (ARI) from running the various methods on simulated noisy datasets. Datasets were created by adding either adding 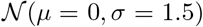 noise (constant layer) or 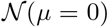 with standard deviation *σ* ∈ (0, 4) (layer with varying amount of noise). We report ARI of the classification at each data point for 100 repetitions.

**Figure S3:**
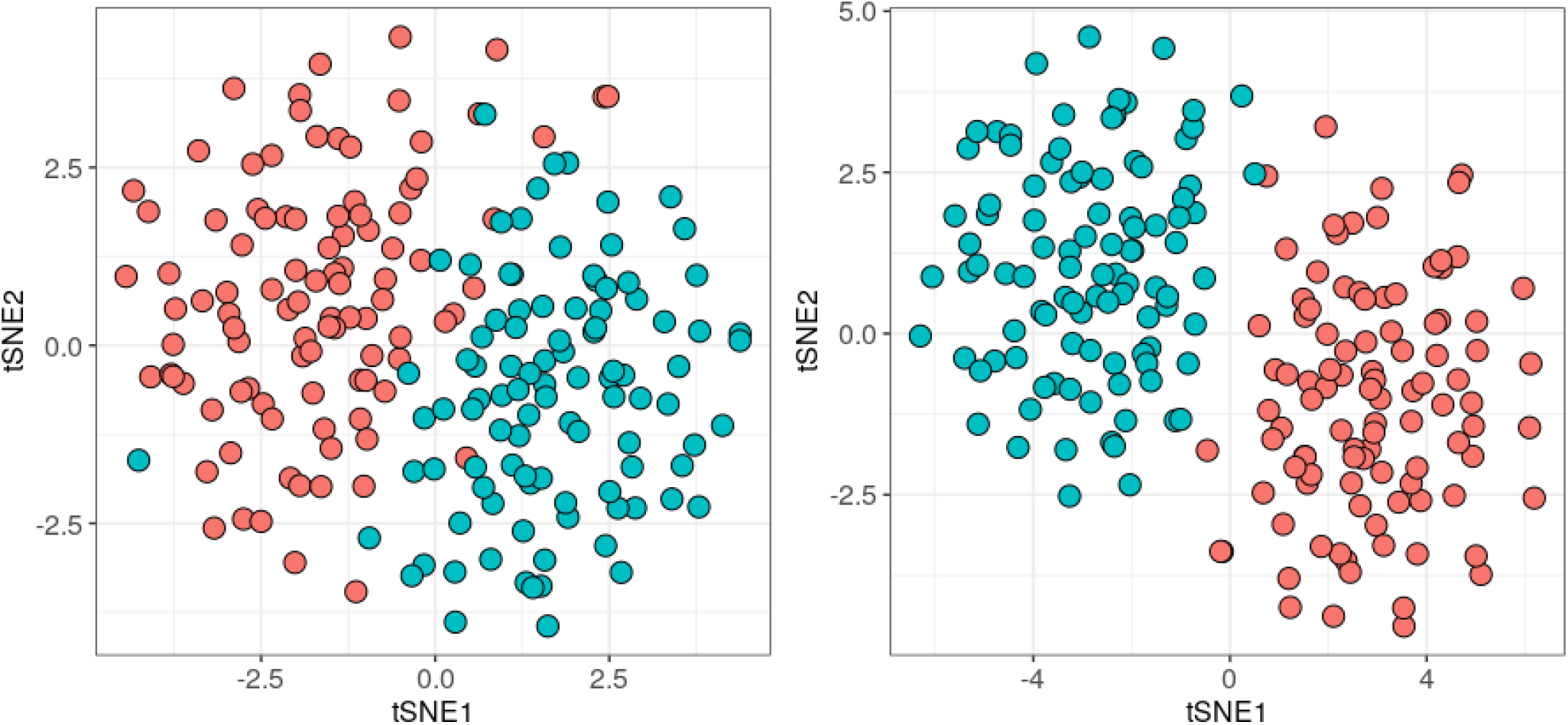
Experimental setup to compare the accuracy of the various methods with varying sample size. Cluster separability in stability simulations show here using tSNE. Data type shown in the left panel contains noise from 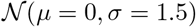 distribution, while the one on the right has noise from 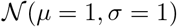 distribution.

**Figure S4:**
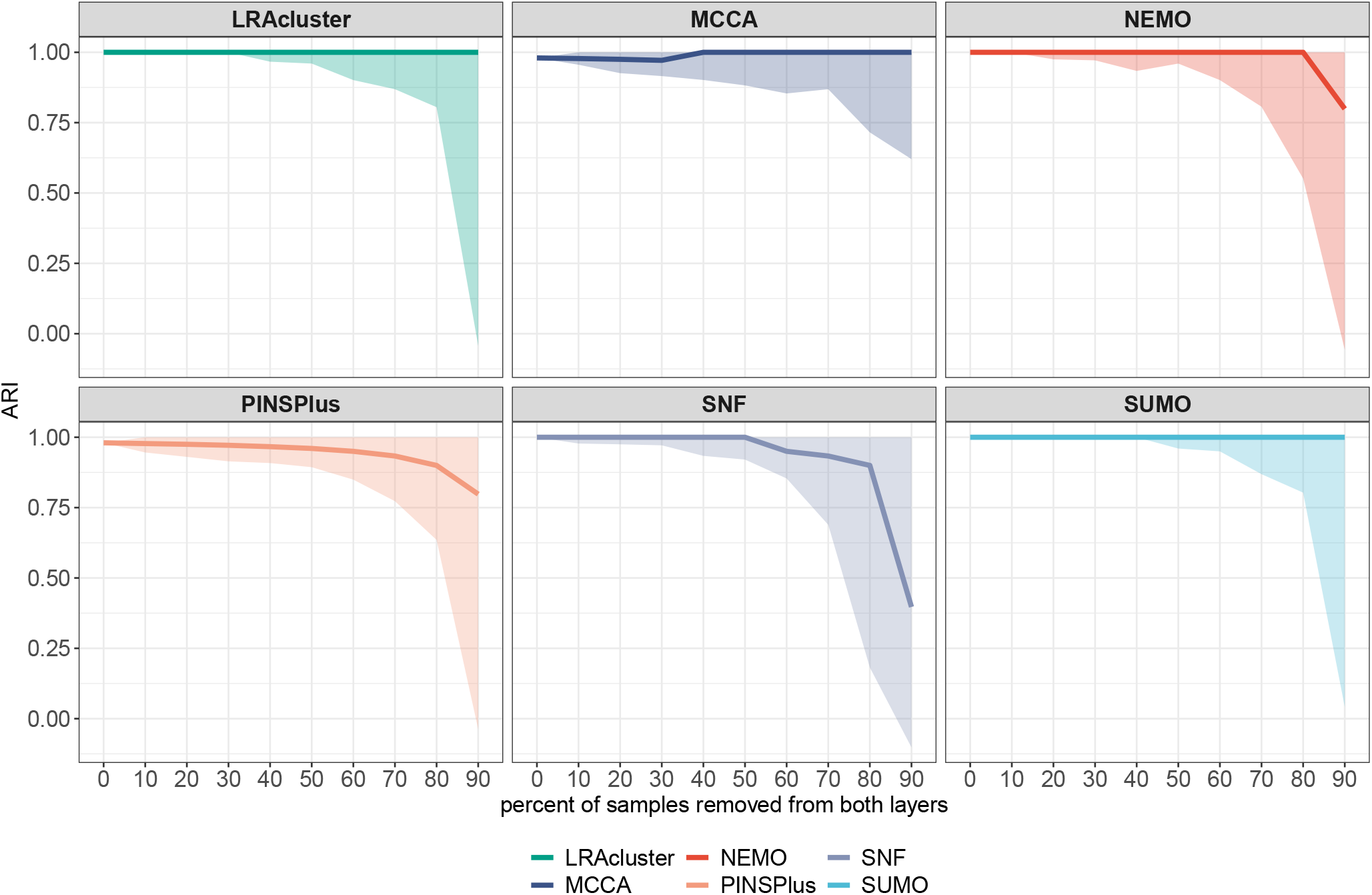
Effect of sample size on the accuracy of the various methods. Datasets were created by removing the same random fraction of samples from both data type. We plot the ARI scores for 100 repetitions at each data point.

**Figure S5:**
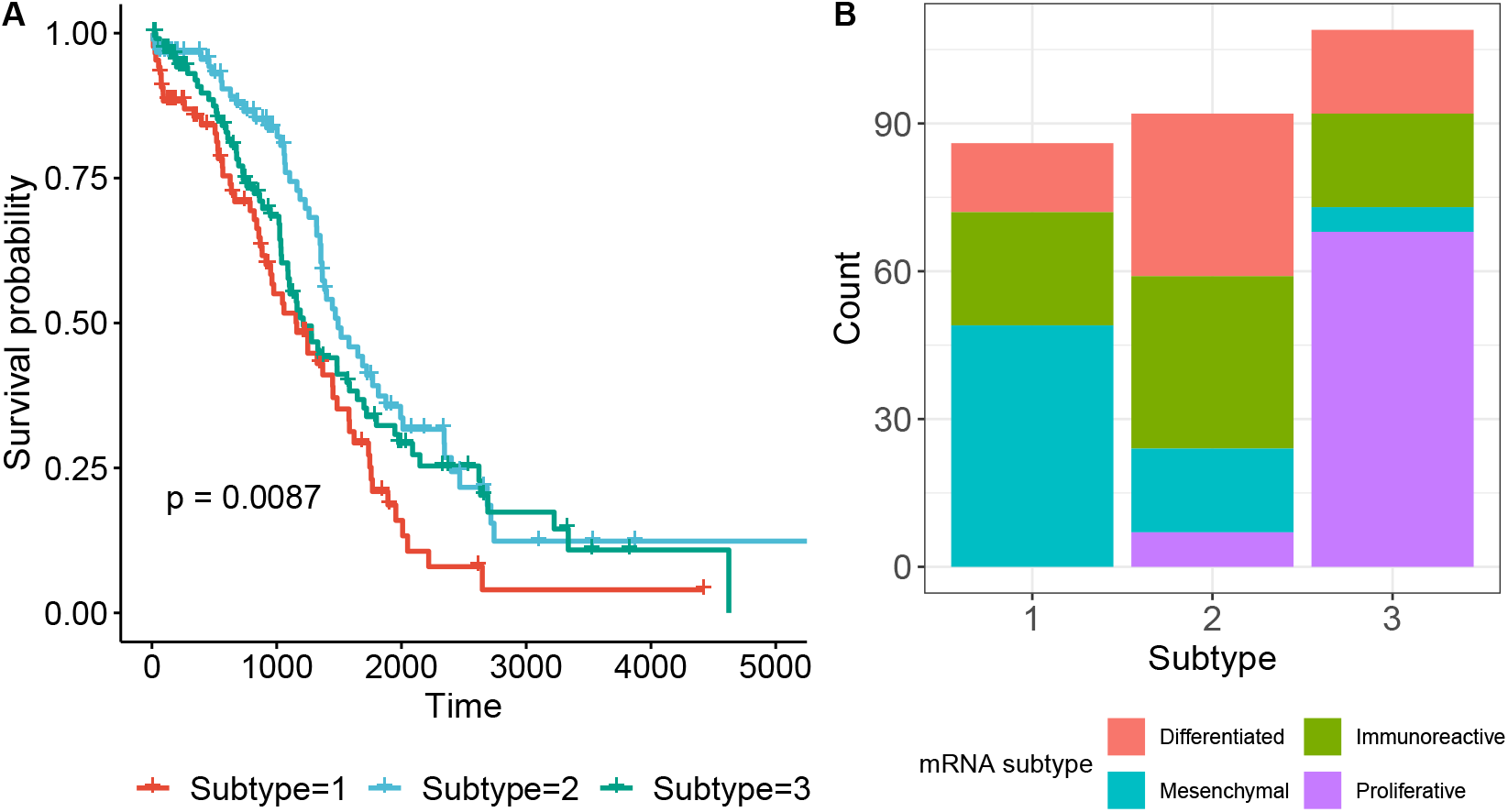
SUMO identifies a subgroup of patients with significant differential survival in the TCGA-OV dataset. (A) shows the KM analysis of the clusters identified by SUMO, and (b) shows the distribution of the patients based on the subtypes identified by using mRNA dataset.

**Figure S6:**
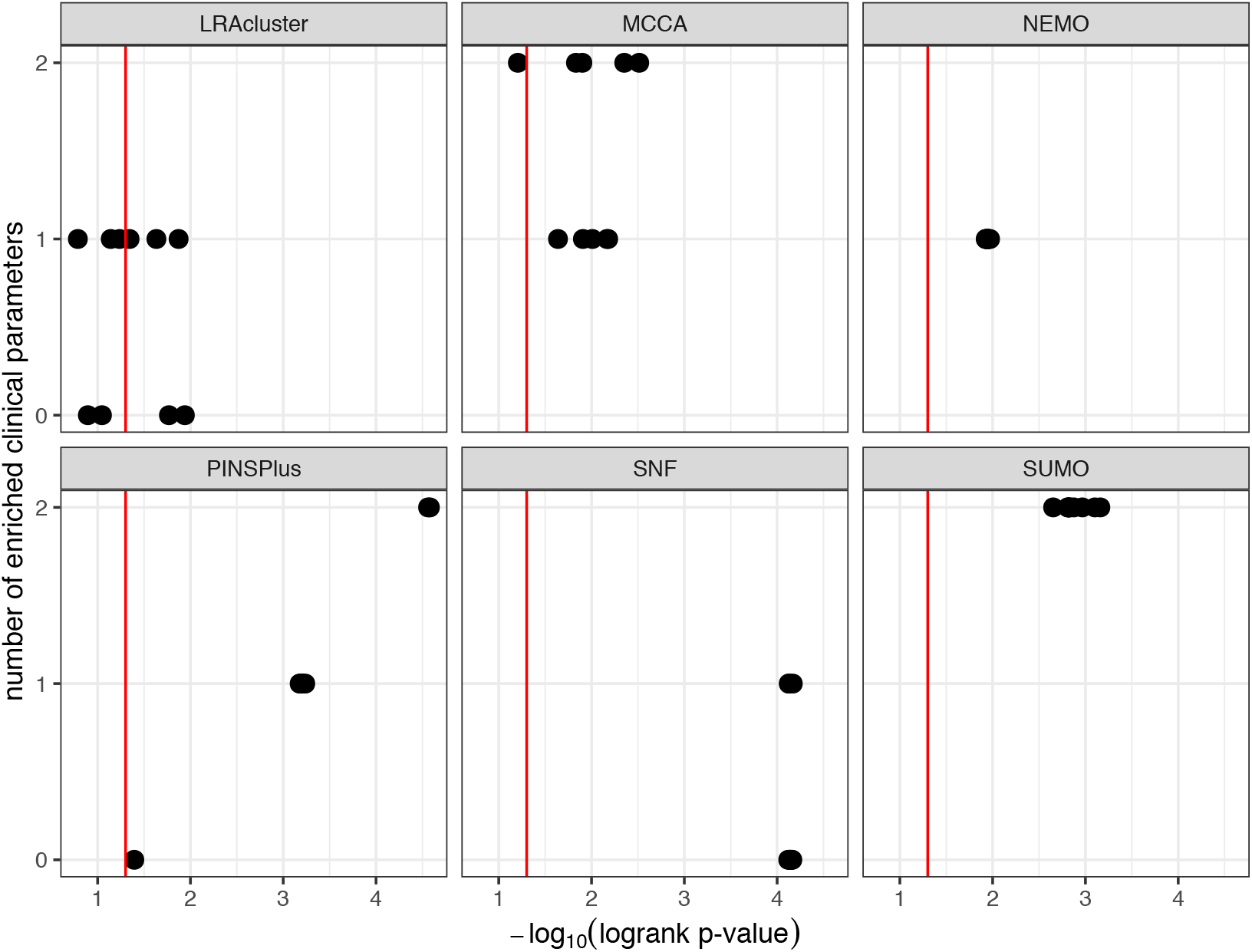
Results of benchmark evaluation using GBM dataset. Vertical line indicates p-value equal 0.05. Each method was run 10 times on this dataset using random seeds as input.

**Figure S7:**
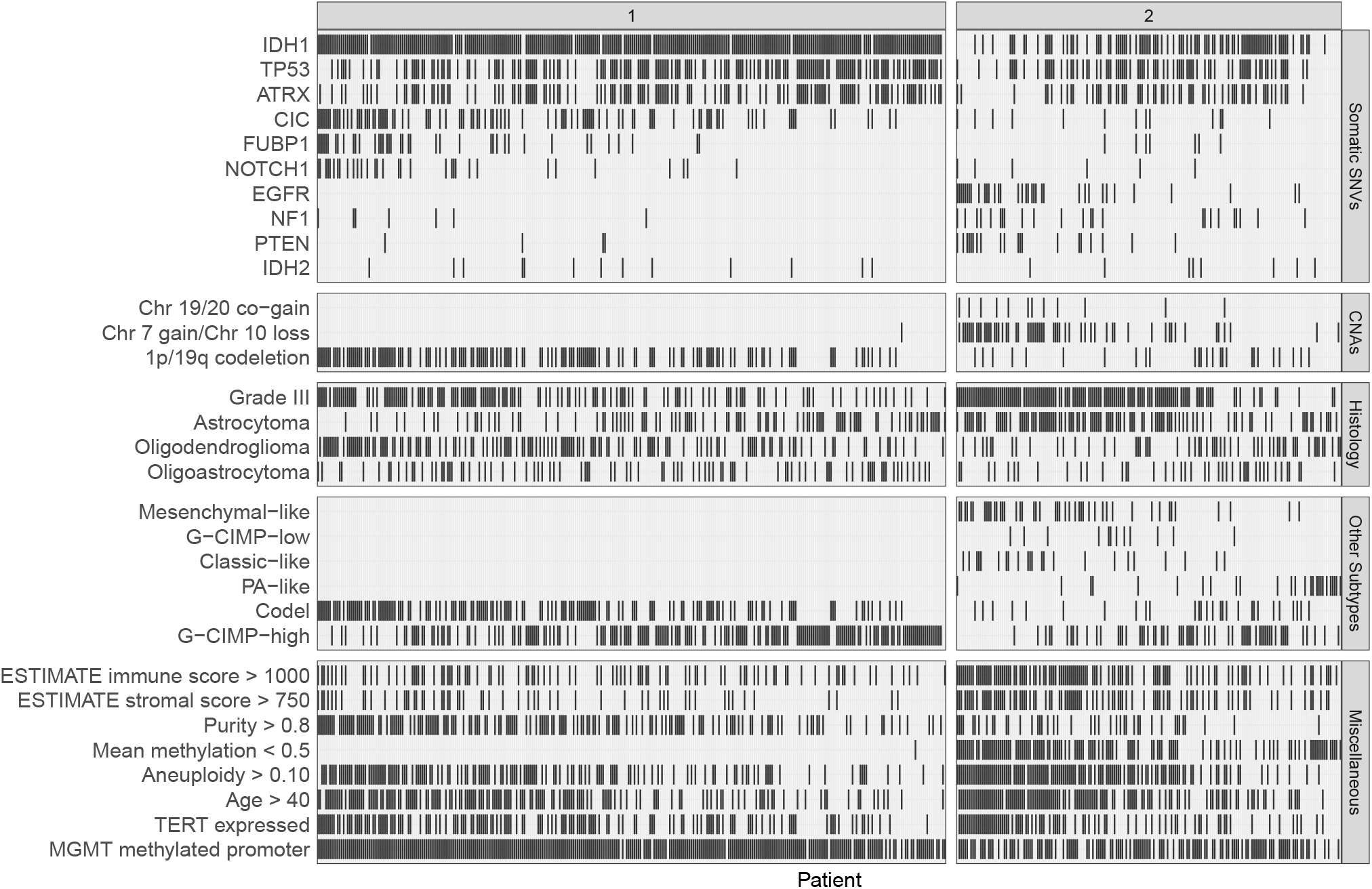
Association of the 2 subtypes identified by SUMO with mutations, clinical phenotypes, and existing supervised classifications.

**Figure S8:**
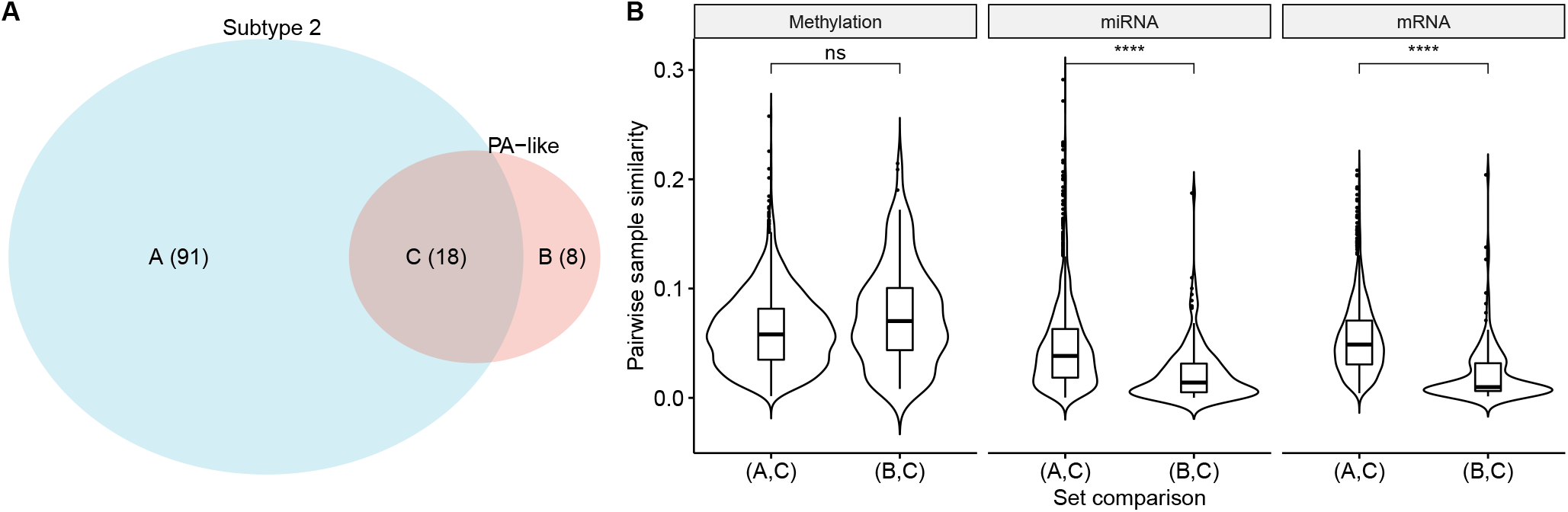
PA-like samples in subtype 2 compared to other samples in subtype 2 and PA-like samples assigned to other subtypes. (A) 18 of the 26 PA-like samples are assigned to Subtype 2 by SUMO. (B) The distribution of pairwise similarity between samples, calculated from euclidean distances. Label A,B,C in the x-axis point to sets of samples in (A). PA-like samples assigned to subtype 2 are more similar to other samples assigned to Subtype 2, when data from mRNA and miRNA are included in the analysis.

**Figure S9:**
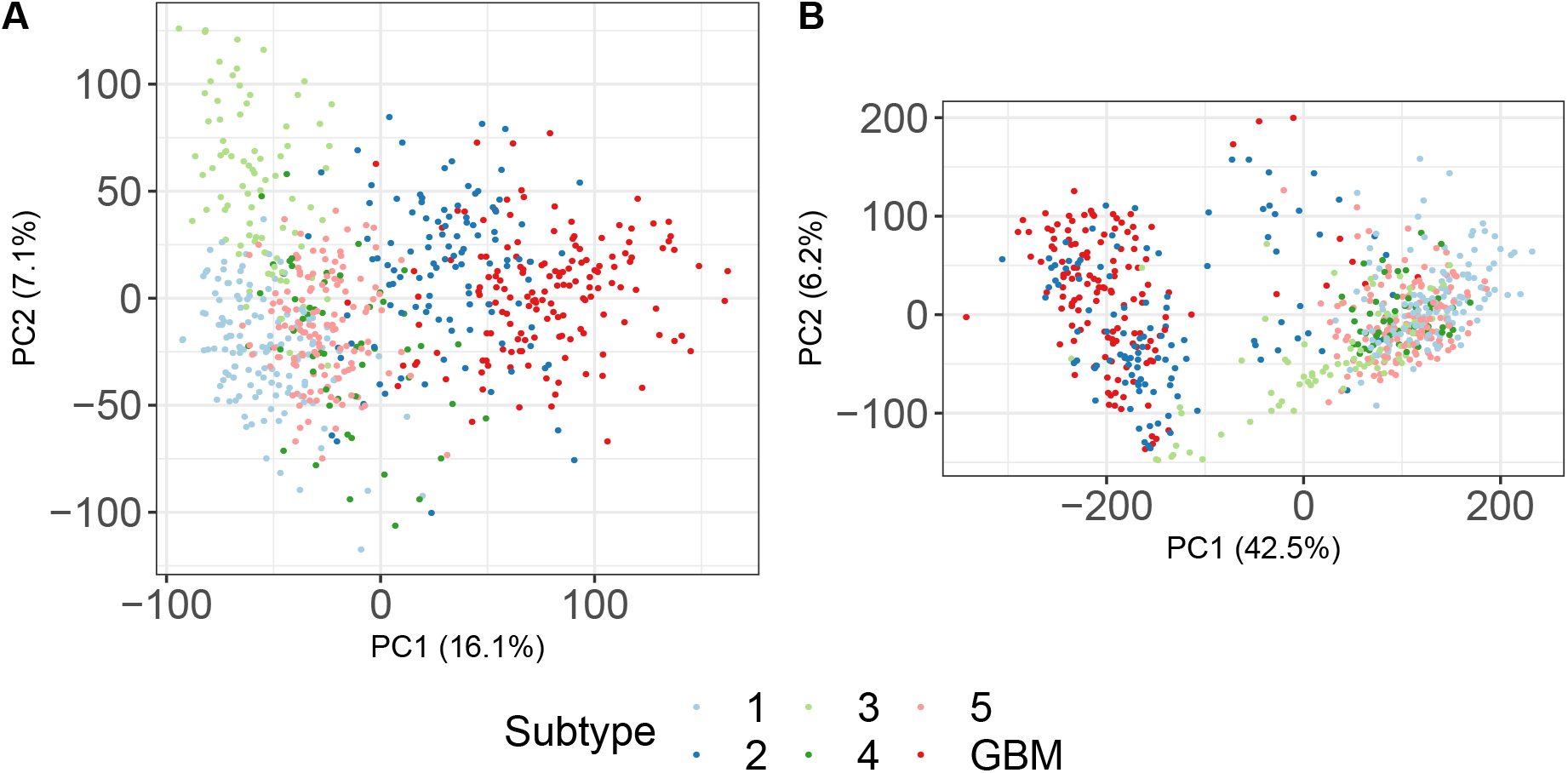
Principal component analysis. We show the top 2 principal components from (A) expression, (B) methylation of TCGA-GBMLGG which shows the similarities between Subtype 2 and GBMs.

**Figure S10:**
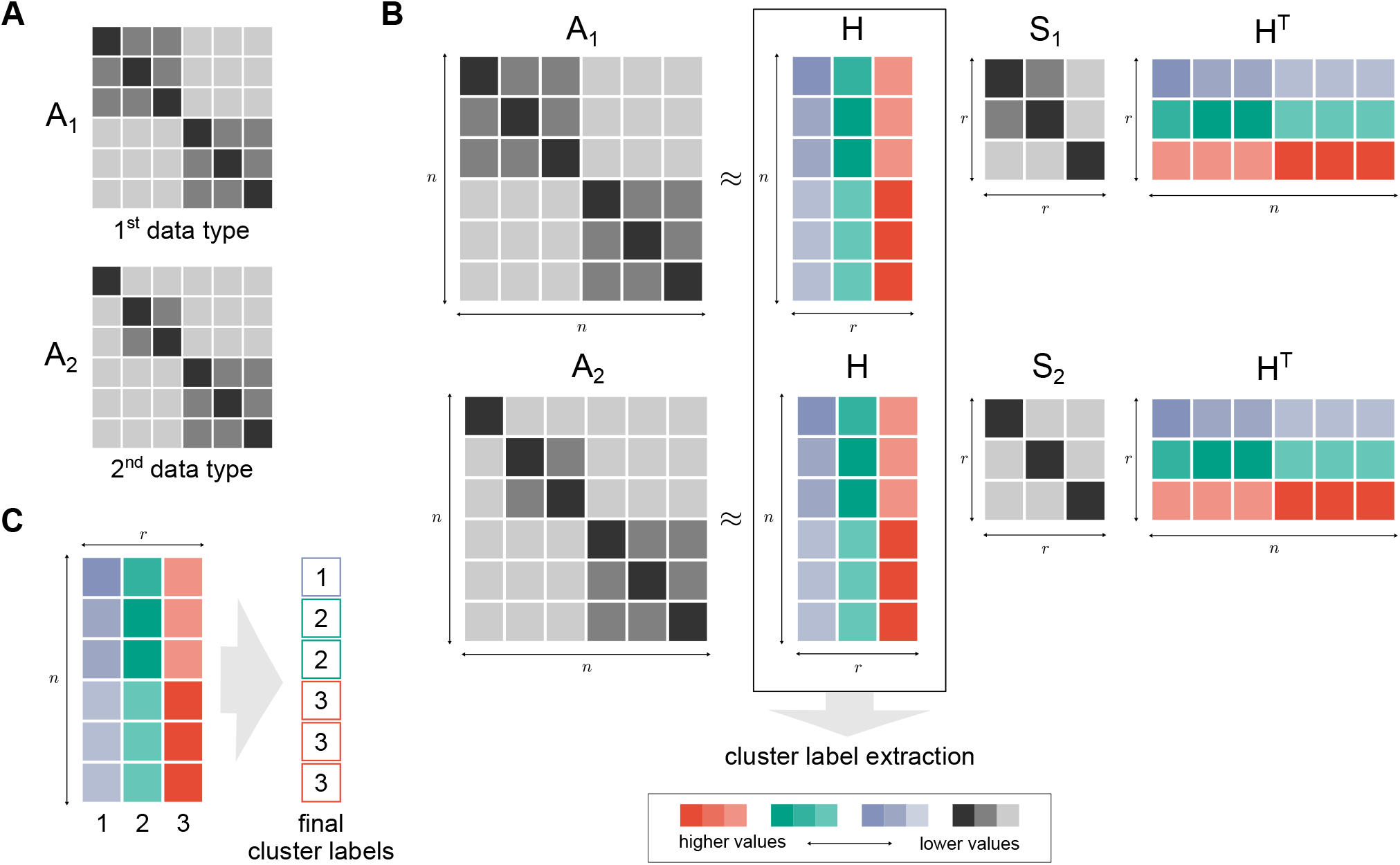
Illustrative description of factorization. (A) Two similarity matrices *A*_1_ and *A*_2_ display complementary sample-sample similarity in both data types. (B) Each similarity matrix is tri-factorized in such a way that *H* matrix is shared across the data types and afterward used for cluster label extraction. Data type specific *S_i_* matrices display relationships between clusters. (C) Final cluster labels for samples are extracted by inspecting columns containing row-wise maximum values of *H* matrix.

**Figure S11:**
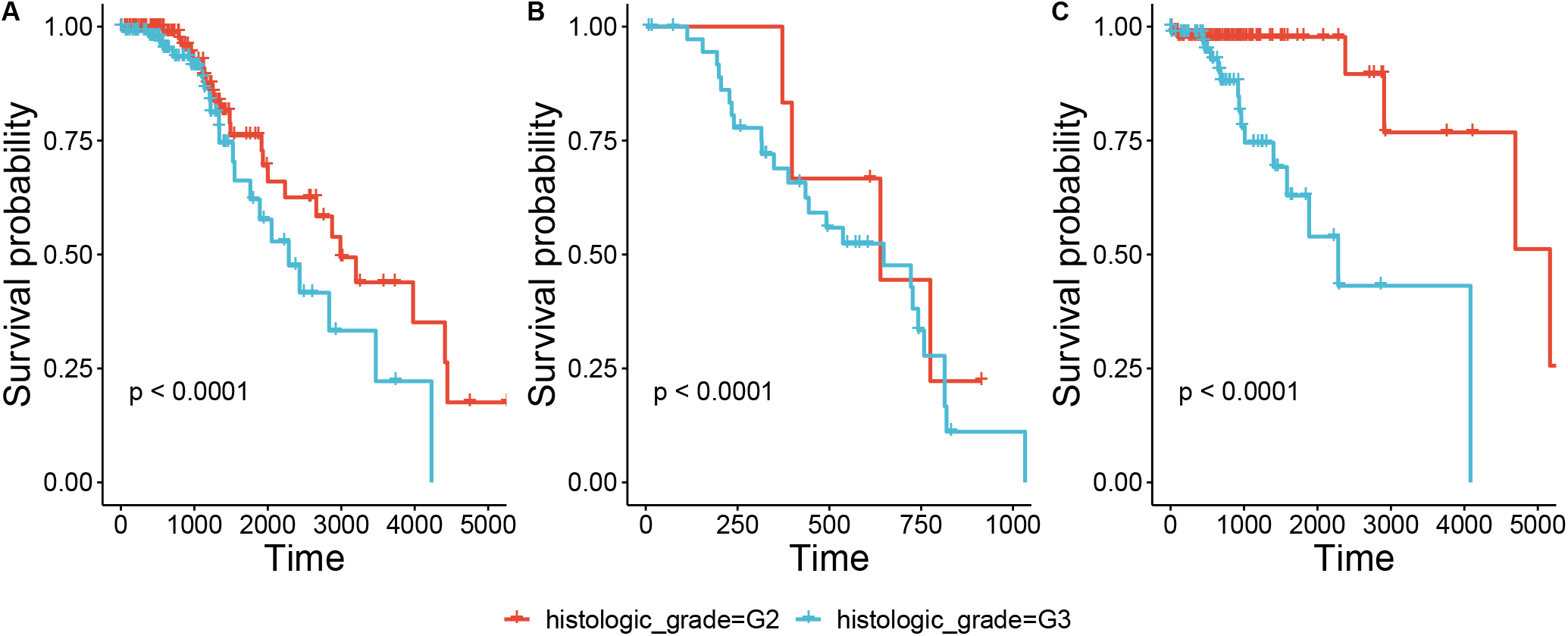
There is a significant difference in survival within the subtypes based on tumor grade. Here we show the samples assigned to (A) G-CIMP-high, (B) Mesenchymal-like, and (C) Codel subtypes and show that KM analysis finds significant differences in survival based on the assigned tumor grade.

**Figure S12:**
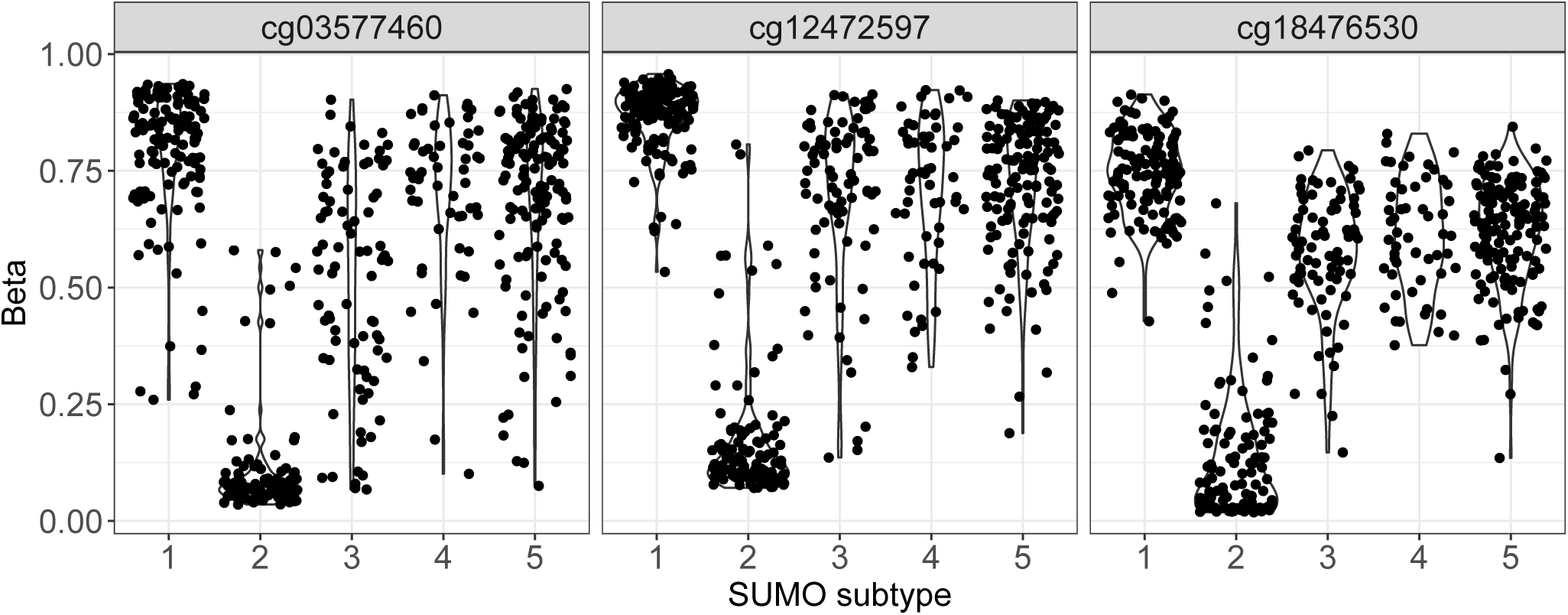
The top three features with the potential to be biomarkers for Subtype 2. Here we show the violin plot of the beta values for the probes with the highest predictive values, as identified on running the interpretation module in SUMO.

**Figure S13:**
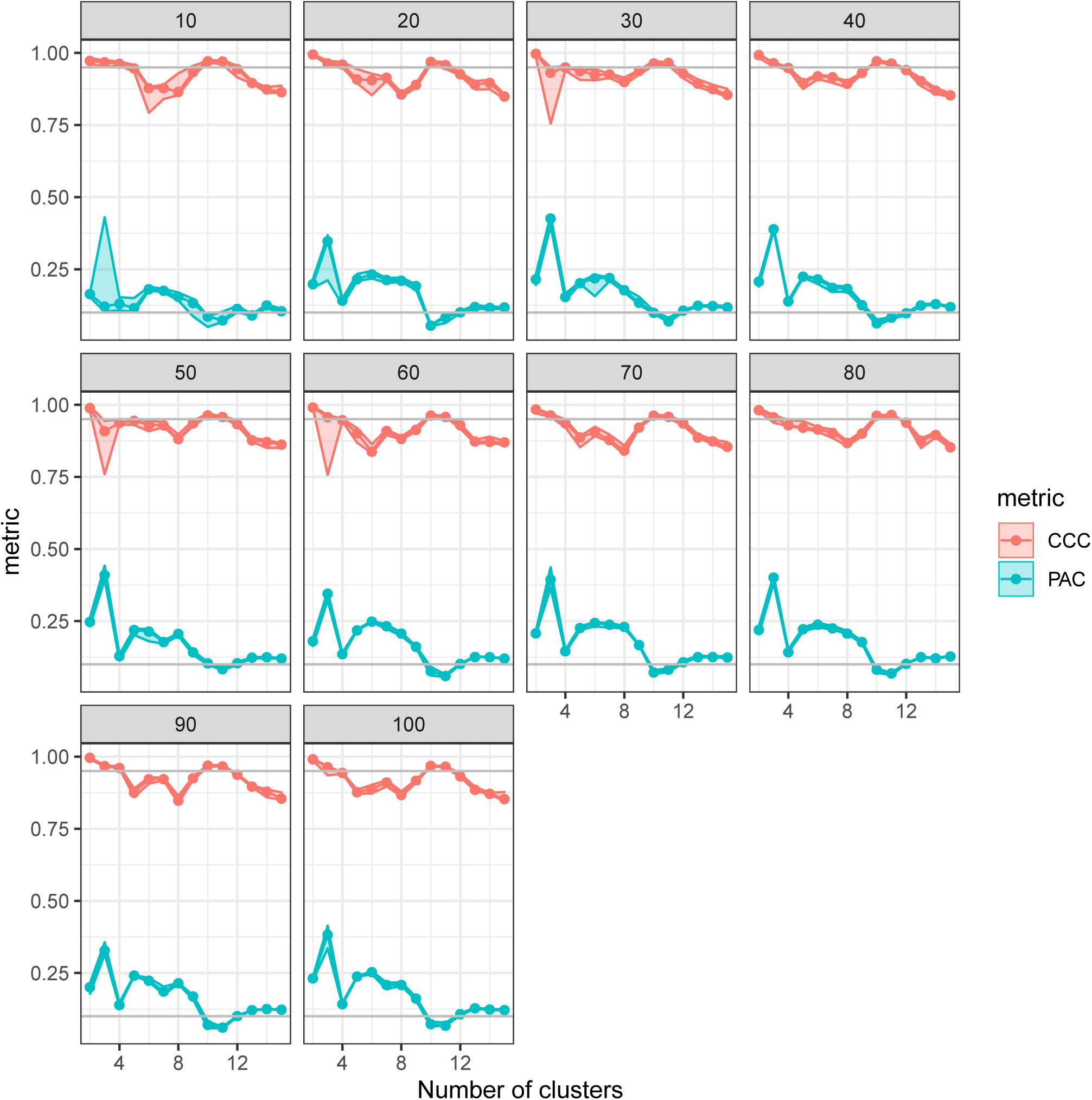
SUMO can identify one of the stable solutions after 20-30 repetitions. Here, each facet shows the PAC and CCC curves (the minimum, median and the maximum value of those metrics are shown for each “number of clusters”) as the number of repetitions of the solver is increased. SUMO identifies either 10 or 11 as the optimal number of clusters when a small number of repetitions are run. As the number of repetitions increase, both 10 and 11 emerge as equally stable solutions. The horizontal lines correspond to values of 0.1 and 0.95.

**Figure S14:**
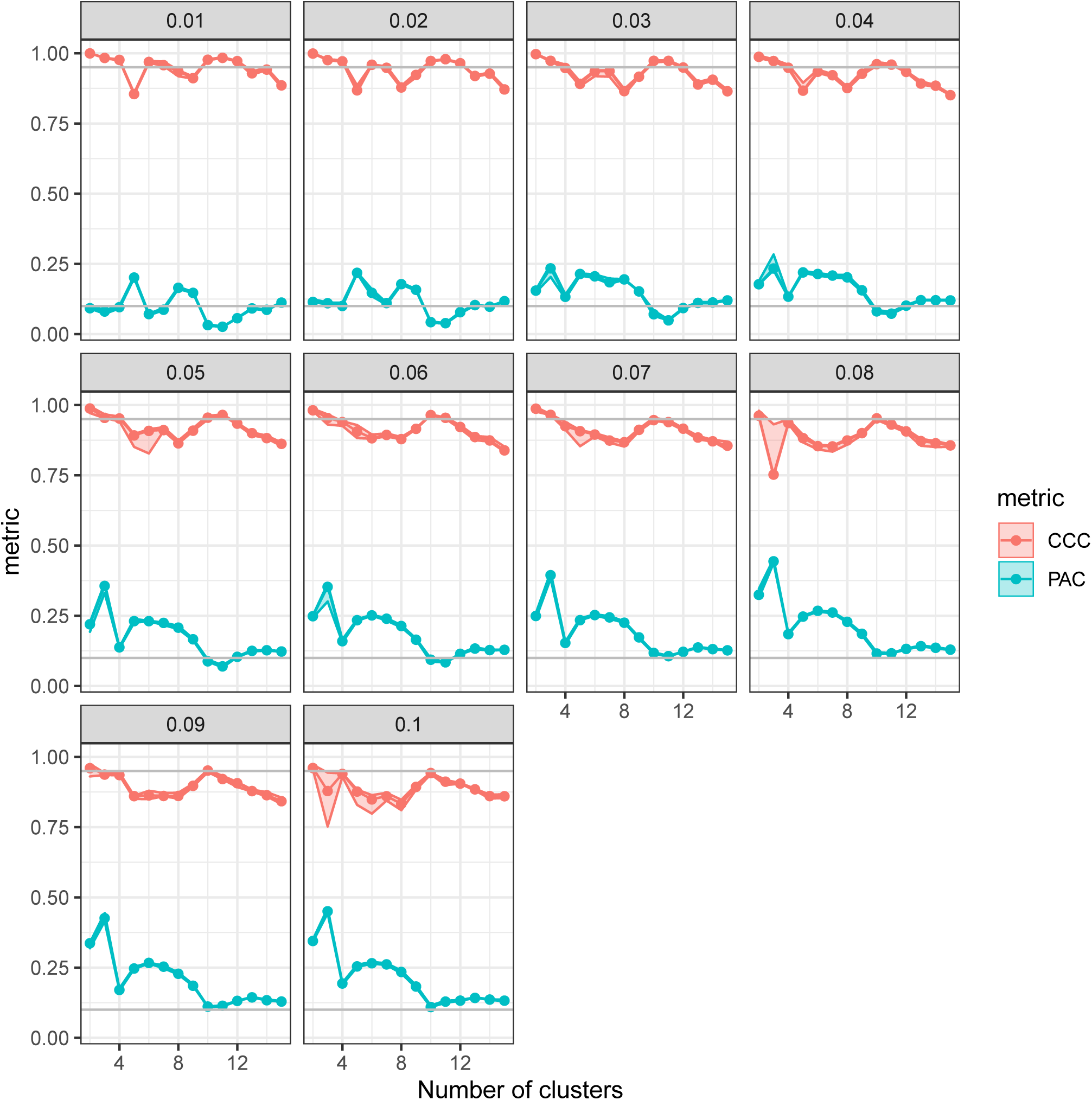
SUMO is stable for a wide fraction of samples that are removed in a single repetition. Here, each facet shows the PAC and CCC curves (the minimum, median and the maximum value of those metrics are shown for each “number of clusters”) as the fraction of samples that are removed in each of repetitions of the solver is varied. SUMO identifies either 10 or 11 as the optimal number of clusters as the fraction is change from 1% to 10% of the samples. The horizontal lines correspond to values of 0.1 and 0.95.

